# Two transcriptional cascades orchestrate cockroach leg regeneration

**DOI:** 10.1101/2023.12.09.570905

**Authors:** Chonghua Ren, Yejie Wen, Shaojuan Zheng, Zheng Zhao, Ethan Yihao Li, Chenjing Zhao, Mingtao Liao, Liang Li, Xiaoshuai Zhang, Suning Liu, Dongwei Yuan, Wei Wang, Jifeng Fei, Sheng Li

## Abstract

The mystery of appendage regeneration has fascinated humans for centuries, while the regulatory mechanisms remain unclear. In this study, a transcriptional landscape of regenerating leg was established in the American cockroach, *Periplaneta americana*, an ideal model for appendage regeneration with remarkable regeneration capacity. Through a large-scale *in vivo* screening, we identified multiple signaling pathways and transcription factors (TFs) controlling leg regeneration. Specifically, *zfh-2* and *bowl,* which have not been previously implicated in appendage regeneration, contributes to blastema proliferation and morphogenesis in two novel transcriptional cascades BMP/JAK-STAT-*zfh-2*-*bab1*/*B-H2/Lim1* and Notch-*drm/bowl*-*bab1*. Notably, *zfh-2* was found working as a direct target of BMP signaling to promote cell proliferation in the blastema. These mechanisms might be conserved in the appendage regeneration of vertebrates from an evolutionary perspective. Overall, our findings reveal that two crucial transcriptional cascades orchestrate distinct cockroach leg regeneration processes, significantly advancing the comprehension of molecular mechanism in appendage regeneration.

## Introduction

“If there were no regeneration, there would be no life” ^1^. Regeneration is an important function that allows organisms to repair damaged or even amputated body parts. The regenerative abilities of animals vary greatly, with some species capable of complete body recovery, like flatworms, and others exhibiting strong regeneration in specific tissues, such as fins in fishes and appendages in arthropods and amphibians. In contrast, mammals typically have limited or no capacity for appendage regeneration. Limb or leg regenerations has been extensively studied due to their vital role in facilitating locomotion and other essential daily activities ^2^. Insects, as the most diverse metazoan taxon, offer a wealth of research opportunities for investigating leg regeneration due to various degrees of regenerative capacity observed across thousands of species. Related research provides an evolutionary perspective view on leg regeneration ^3^. Similar to other invertebrate and vertebrate species ^4–6^, insect leg regeneration processes can be divided into three distinct stages: wound healing, blastema formation, and morphogenesis (including regenerative growth, differentiation, and patterning). The blastema forms at the wound site and generates various cell types that contribute to leg regeneration. These cells then undergo proliferation, and the newly grown tissue re-establishes proper patterning to restore leg functionality ^7^. Although the mechanisms behind appendage regeneration have fascinated humans for centuries, they still remain largely unclear in most cases ^8^.

To ensure proper replacement of the damaged leg, precise orchestration of the transcriptional landscape is required. Dynamic regeneration processes involve sequential waves of gene expression as transcriptional cascade mechanisms through transcription factors (TFs), and step-by-step regulation of TFs and related signaling pathways form regulatory networks ^9^. Several signaling pathways, such as <u>J</u>un <u>N</u>-terminal <u>k</u>inase (JNK) ^10^, Wingless(Wnt/Wg) ^11,12^, Hedgehog (Hh) ^12,13^, Decapentaplagic (Dpp) ^12,14^, <u>e</u>pithelial <u>g</u>rowth factor <u>r</u>eceptor (EGFR) ^15^, and <u>Ja</u>nus <u>k</u>inase-<u>s</u>ignal transducer and <u>a</u>ctivator of transcription (JAK-STAT) ^16,17^, have been found to play critical roles in leg blastema formation and morphogenesis in crickets, fruit flies, and other insects. Several TFs in these signaling pathways, such as *Stat*, *mothers against Dpp* (*Mad*, also named *Smad1*), and *cubitus interruptus* (*ci*), have been highlighted. In addition, two other TFs, *distal-less* (*Dll*) and *dachshund* (*dac*), have been demonstrated to play key roles in regulating proximodistal (P/D) pattern formation under the control of the Dpp and Wnt/Wg signaling pathways ^18,19^. However, systematic research on the hierarchical transcriptional cascades that control blastema formation/proliferation and/or morphogenesis in a given insect species is absent ^7^.

Cockroaches (Blattodea) are attractive objects in regeneration research because their leg regeneration ability is top-ranked among Insecta ^7^. In cockroaches, many regenerative hypotheses, such as intercalary regeneration and proximal-distal axis, have been established, and several conceptual models, such as the gradient model, polar coordinate model and boundary model, have been proposed ^20^. The American cockroach, *Periplaneta americana*, is a raw material in traditional Chinese medicine, and substances extracted from *P. americana* (including the prescribed drug *Kangfuxin Solution*) are used in the clinic for tissue repair and to treat ulcerative diseases ^21^. There may be a connection between the powerful leg regeneration ability in this species called as “Xiao qiang (Little mighty)” and its remarkable clinical utility ^22^. Our previous study showed that *Dpp* and *Mad* are indispensable for leg regeneration in this cockroach species ^14^. Nevertheless, the transcriptional regulation mechanisms in cockroach leg regeneration remains unknown.

In this study, the temporal transcriptional landscape of regenerating *P. americana* legs was established using comparative transcriptomic analysis upon detailed morphological recording. Two transcriptional cascades based on TFs, *zfh-2* and *bowl*, were found to predominantly control blastema cell proliferation and/or morphogenesis. We further show that these two novel transcriptional cascades lie in traditional signaling pathways (e.g. Dpp, JAK-STAT, and Notch) that form a network for regulating different leg regeneration processes. Significantly, the discoveries in this ideal insect model provide broad insights into transcriptional regulation of appendage regeneration from an evolutionary perspective.

## Results

### Morphological Profiles of Leg Regeneration

As shown in our previous work, the trochanter plays key roles in leg regeneration ^14^; thus, in this work, all amputations were performed at the trochanter-femur joint. New legs complete in shape and function were regenerated after one molting (∼85% of the contralateral leg) (Fig. 1A and S1A). To profile different regeneration processes, multiple time points 6 h, 12 h, 1 d, 3 d, 5 d, 7 d, and 10 d post amputation (pa) were selected for three types of morphological examinations. We first performed micro<u>c</u>omputed tomography (μ-CT) scanning to record regenerating microstructural objects in great details as a three-dimensional (3D) model. The regenerating processes were observed from four (front to back, left to right) different directions (Fig.1B and S1B). To further understand the regeneration process at the cellular level, 5-ethynyl-2’-deoxyuridine (EdU) labeling and histochemical hematoxylin-eosin (HE) staining were performed to detect tissue change and cell activity (Fig. 1C and S1C). At the beginning of regeneration, a clot forms at the amputation site. Hemocytes aggregate in the wound region to eliminate the inflammatory response ^23^, epidermal cells migrate over the wound surface and epidermal continuity is restored underneath the scab (∼6 hpa-1 dpa). The cells in the wound region begin to regain proliferation ability by 1 dpa, blastema is formed in the wounded trochanter (∼3 dpa), and elongates accompanied by the formation of original shape of the new leg (∼5 dpa). Epidermal cells of the blastema and primordia of regenerating tissues have high proliferative activity by 3-5 dpa. After this point, regenerating tissues continue to grow and fold inside of the old coxa until inflation and extension after molting (∼7-10 dpa) (Movie.S1). The cell proliferation peaks at 7 dpa and is decreased by 10 dpa. In summary, our comprehensive morphological characterizations, based on μ-CT, EdU, and HE analyses, clearly defined three distinct regeneration phases that have been shown in vertebrate appendage regeneration, wound healing (∼6 hpa-1 dpa, dedifferentiation), blastema formation (∼3-5 dpa, proliferation), and morphogenesis (∼5-10 dpa, proliferation and differentiation) (Fig. 1D). The morphological profiles lay a solid foundation for our subsequent transcriptional profile research.

**Fig. 1.**
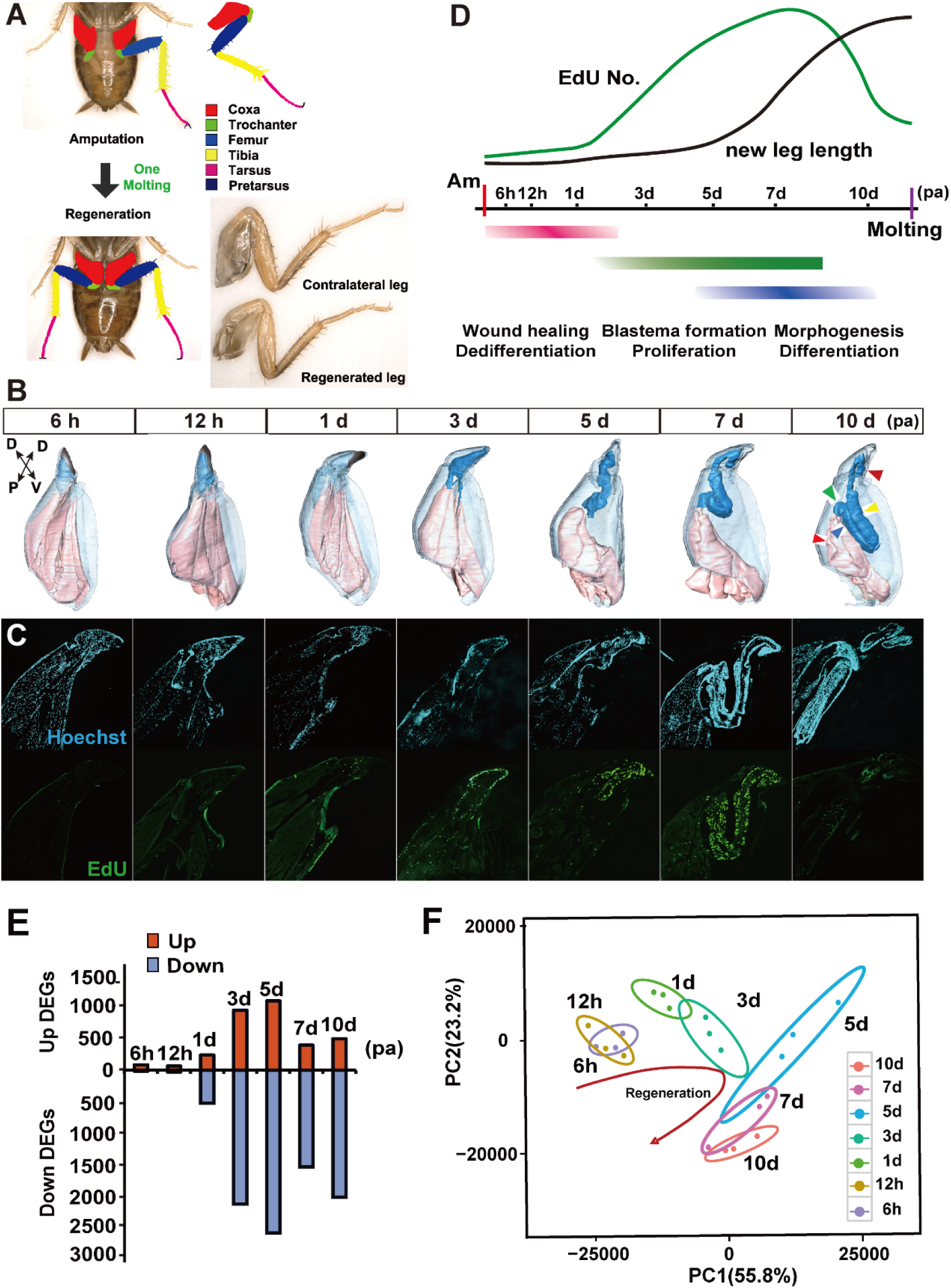
Morphological and transcriptional profiles of leg regeneration in American cockroach. (A). The diagram of typical and regenerated legs. Six different colors represented different segments of legs. (B). 3D model of the morphological profiles of regenerating legs made by μ-CT. Front and back directions were adopted to show the regenerating microstructural objects. Black regions indicated wound scar, blue regions indicated new regenerated tissues, and pink regions indicated the remaining tissues in the coxa; P-D and D-V indicated the proximal-distal and dorsal-ventral axis, respectively. Different segments of regenerated legs were pointed with the corresponding colors same to 1A. (C). EdU staining of proliferated cells during 24 h period before harvest. Cell nucleus were stained with Hoechst dye. (D). Diagram of leg regeneration processes and trends of the number of EdU positive cells and the length of new regenerated legs. (E). Statistics of the number of differentially expressed genes at different time points. Red histograms indicate upregulated genes, blue histograms indicate downregulated genes. (F). The profiles of different expressed genes at 7 time points during regeneration analyzed by the PCA.

### Transcriptional Profiles Reveal Dynamic Gene Expression Patterns

To uncover the transcriptional dynamics of regenerating legs, the distal coxa and the whole trochanter of the one-side amputated (AM) legs at the same time points described above were harvested and subjected to transcriptome sequencing. To rule out the influence of leg homeostasis on gene expression, corresponding distal coxa and trochanter of the contralateral (CL) legs that had not undergone amputation were used for comparison (Fig. S2A). The differentially expressed genes (DEGs) at every time point between the two groups are preferred indexes for evaluating the transcriptional profiles of regeneration (Table. S1). At the early wound healing stage, there were few DEGs at 6 hpa and 12 hpa, with a slight increase at 1 dpa. In the middle blastema formation stage, a significant increase in the number of DEGs occurred at 3 dpa and peaked by 5 dpa. After that, the number of DEGs decreased in the late morphogenesis stage at 7 dpa and 10 dpa (Fig. 1E). In general, there are more downregulated genes than upregulated ones from 1 dpa to 10 dpa. This observation suggests that the downregulated genes may also play important roles in leg regeneration, in addition to the expected upregulated genes. Collectively, an obvious trend in the expression patterns of DEGs across the seven regenerative time points was observed by principal component analysis (PCA) (Fig. 1F), aligning with the observed morphological changes. We then performed hierarchical clustering analysis to systematically compare the DEGs that appeared at each of the seven time points, and nine main clusters were identified when a parameter of fifteen was set. Gene Ontology (GO) enrichment analyses were performed for both up or down regulated genes, and several processes including structural constituent of cuticle, oxidoreductase activity, RNA binding, and ATP binding were found (Fig. S2B). In addition, several canonical signaling pathways associated with development and regeneration were found in these clusters, such as transforming <u>g</u>rowth factor-<u>β</u> (TGF-β) that includes <u>b</u>one <u>m</u>orphogenetic <u>p</u>rotein (BMP, a major subfamily of TGF-β family) pathway, Hippo, Wnt/Wg, Hh, Notch, InR-FoxO, and ECM (could be degraded by matrix metalloproteinases, MMPs) ^24,25^ (Fig. S2B), implying that these signaling pathways might be involved in cockroach leg regeneration. In brief, we devised a protocol to isolate distinct DEGs at multiple time points and extract their expression profiles using RNA-seq, revealing dramatic transcriptional transitions during cockroach leg regeneration, especially at the stages of blastema formation and morphogenesis.

### Multiple Signaling Pathways Are Essential for Blastema Proliferation and Morphogenesis

To verify the necessity of the essential signaling pathways in cockroach leg regeneration, all seven pathways enriched above and five not enriched but considered strong candidates for functions on regeneration (JAK-STAT, EGFR, fibroblast <u>g</u>rowth factor receptor (FGFR), <u>J</u>un-<u>N</u>-terminal <u>k</u>inase (JNK), and <u>gr</u>ainy <u>h</u>ead (Grh)) were selected for functional verification. First, five signaling pathways, with exact ligand, receptor, and TF compositions, were selected for *in vivo* RNAi treatments (Fig. 2A-2C and S3). Compared with the ds*Mock* control group (Fig. S3), almost no new tissue was regenerated in the groups with knockdown of *Unpaired* (*Upd*), *Domeless* (*Dome*) and *Stat* (an ortholog of mammalian *Stat3/5* and *Drosophila Stat92E*) in the JAK-STAT pathway (Fig. 2A and 2B). The knockdown of *Dpp*, *Screw* (*Scw*), *Thickveins* (*Tkv*), *Saxophone* (*Sax*), *Mad* and *Smad4* (*Medea*, *Med*) in the BMP pathway ^26^ significantly disrupted the entire regeneration process (Fig. 2A and 2B), which is consistent with our previous finding ^14^. However, silencing of the other two ligands in the BMP pathway, *Glass-bottom boat* (*Gbb*) and *Maverick* (*Mav*), produced no obvious phenotypic defects (Fig. S3). The RNAi screening indicates that BMP signaling contributes to leg regeneration through the ligands Scw and Dpp rather than Gbb or Mav. When *Delta* (*Dl*), *Notch* (*N*), and *Suppressor of Hairless* (*Su(H)*) were knocked down, the regenerated legs displayed significantly reduced length and distorted patterning. The joint sites among the femur, tibia and tarsus exhibited a swelling phenotype, rendering them non-functional (Fig. 2A and 2B). The Wnt/Wg pathway demonstrated a relatively weak impact on leg regeneration. In case of *Wg* or *Arrow (Arr)* knockdown, regenerated legs exhibited morphological abnormalities in femur and tibia patterning, and tarsus regeneration was markedly impeded. Conversely, no noticeable defect was observed after *Pan/TCF* RNAi (Fig. 2A and 2B). For the Hh pathway, the roles of *patched* (*ptc*) and *ci* were mainly reflected in the size of legs, segmentation of the tarsus was abolished under *ci* RNAi, while no obvious defects were found after *Hh* silencing (Fig. 2A and 2B). These results suggest that the JAK-STAT and BMP signaling pathways control blastema cell proliferation, while Notch, Wnt/Wg and Hedgehog signaling pathways regulate morphogenesis. This conjecture was confirmed by monitoring cell proliferation after inhibiting the TFs *Stat*, *Mad*, *Su(H)*, *Pan*, and *ci* in their respective pathways. *Stat* and *Mad* RNAi significantly reduced the number of mitotically active cells in the blastema detected by EdU and phospho-histone H3S10 (PH3) labeling at 3 dpa (Fig. 2D and 2D’). Nevertheless, no significant decreases of EdU or PH3 levels were found after *Su(H)*, *Pan*, and *ci* knockdown treatments at 3 dpa (Fig. 2D and 2D’). Then, the potential functions of seven other signaling pathways for leg regeneration were verified (Fig. S3). In addition to TGF-β signaling, two other growth factor-associated signaling pathways, EGFR and FGFR, were disrupted. The legs regenerated after silencing *vein* (*vn*) and *Egfr* in EGFR signaling were smaller in size (Fig. S3). Neither knockdown of the ligand *branchless* (*Bnl*) nor the receptor *heartless* (*htl*) in the FGFR pathway resulted in obvious phenotypic defects (Fig. S3). Notably, knockdown of *Yorkie* (*Yki*, *YAP*/*TAZ* in mammals) and *Scalloped* (*Sd*, *TEAD1-4* in mammals) in the Hippo pathway severely disrupted leg regeneration, and the ds*Yki-*treated animals failed to molt (Fig. S3). Grh signaling was also tested, as it has been reported to be essential for wound-dependent epidermal barrier regeneration in *Drosophila* ^27^. A smaller leg was regenerated when *Grh* but not *stit/Cad96Ca* was knocked down. In addition, the JNK (*eiger* (*egr*), *basket* (*bsk*), *Jun-related antigen* (*Jra*), and *kayak* (*kay*)), ECM (*Mmp1*, *Mmp2*), and InR-FoxO (*InR*, *FoxO, PI3K,* and *Tor*) pathways were found to be dispensable for leg regeneration. These results indicate that cockroach leg regeneration is orchestrated by multiple signaling pathways, including JAK-STAT, BMP, and Hippo for blastema proliferation, and Notch, Wnt/Wg, and Hh for morphogenesis. Notably, the TFs in these pathways play key roles in regeneration.

**Fig. 2.**
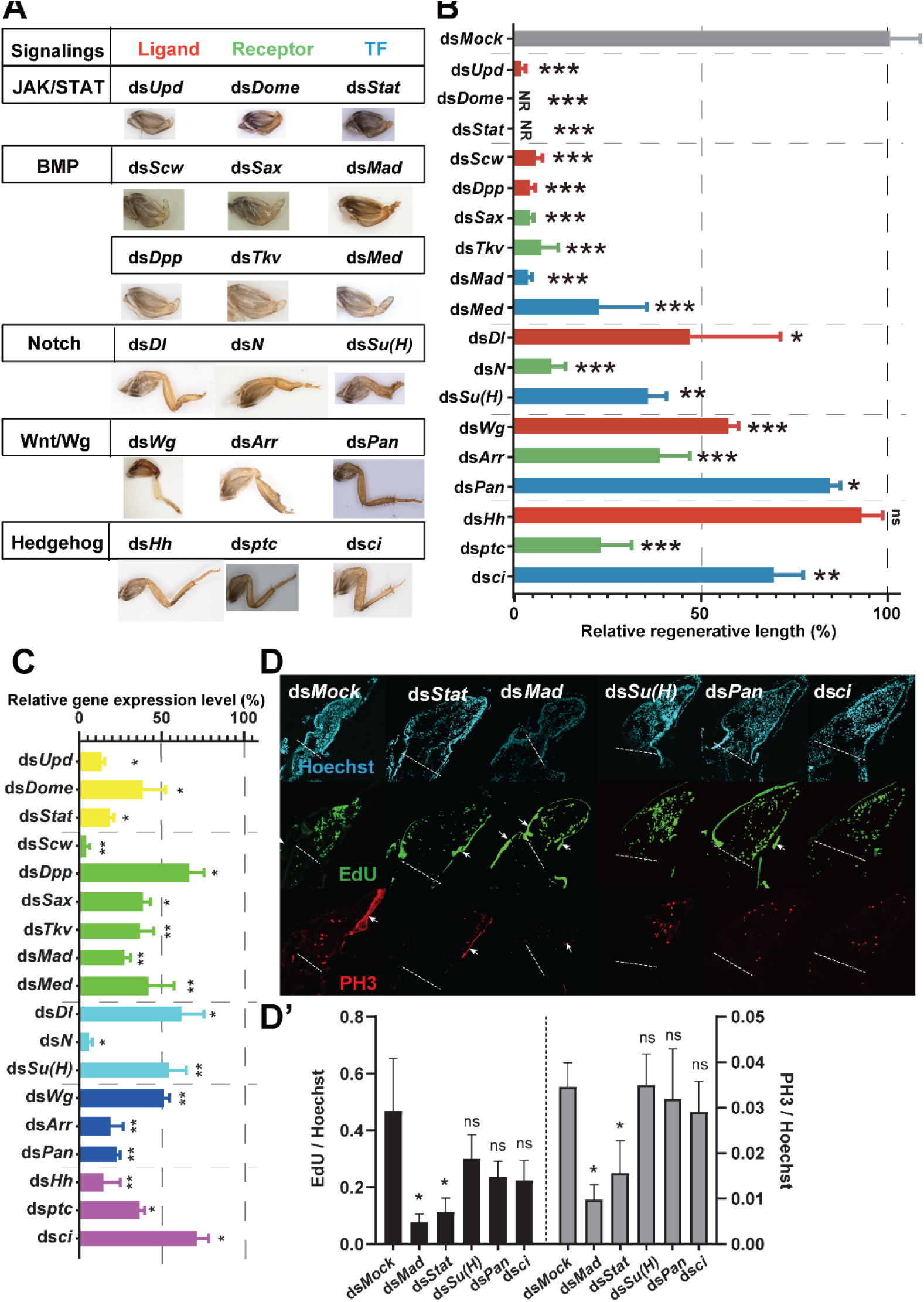
Multiple pathways are essential for blastema cell proliferation and morphogenesis. (A). The key components ligands, receptors, and TFs in five signaling pathways JAK-STAT, BMP, Notch, Wnt/Wg, and Hh were knocked down and their relative regenerative phenotypes were captured. (B). Measurement of the relative regenerative length after RNAi treatments of genes corresponding to them in (A). NR: No visible regeneration found. n=3. (C). The RNAi efficiency of these genes were detected by qRT‒PCR. n=3. (D). EdU and PH3 staining of proliferation positive cells. White arrows indicate the spontaneous fluorescence signal of exoskeleton. (D’). The statistics of the percentages of EdU and PH3 positive cells. dsRNAs were injected at 0 dpa and the samples were harvested at 3 dpa, the EdU was injected at 6 h before harvest. n=3. The significance of differences was analyzed by two-tailed Student’s *t test*. *: *P* <0.05, **: *P* <0.01, ***: *P* <0.001, “ns” stands for no significant difference.

### Large-Scale RNAi Screening for TFs Controlling Leg Regeneration

Focusing on the effect of transcriptional regulation on precise leg regeneration, large numbers of differentially expressed TFs were found in addition to the TFs in the signaling pathways mentioned above (Fig. 3A). The *P. americana* genome annotation ^14^ includes 294 genes homologous to trusted TFs in *Drosophila* (475 genes, Flybase.org). Among these TFs, 178 were found in our RNA-Seq data, and 140 were upregulated in the regenerating legs; and the top 65 TFs with relatively high expression levels were selected for further *in vivo* RNAi screening (Fig. 3A). These 65 TFs can be grouped into 19 classes based on their DNA-binding domains (DBDs). The largest class is the C2H2 zinc finger class with ten (15.4%) TFs, followed by the basic helix-loop-helix class and homeobox classes, each with nine (13.8%) TFs (Fig. 3B). The temporal relative expression profiles of these 65 TF genes throughout the regeneration process are shown in a heatmap (Fig. 3C). The seven TFs (*kay*, *Stat*, *Grh*, *Med*, *Mad*, *Sd*, and *Ci*) that have already been studied as part of the signaling pathways (Fig. 2A) are labeled in green (Fig. 3C). The remaining 58 TFs were subjected to RNAi screening *in vivo*.

**Fig. 3.**
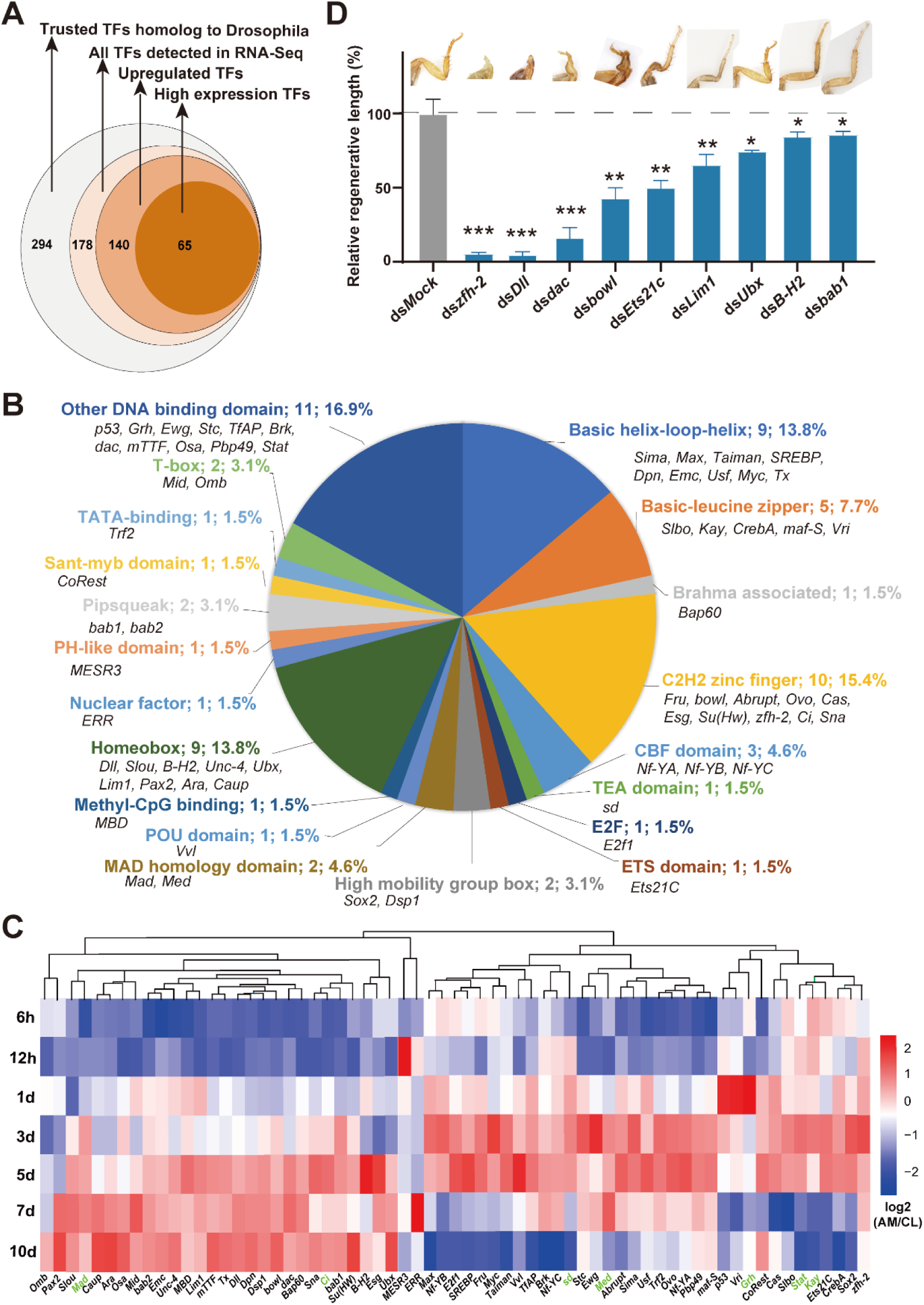
Large-scale screening for TFs controlling leg regeneration. (A). Schematic description of the regenerative RNA-seq gating analysis of TFs. (B). Pie analysis of the selected 65 TFs, and they are divided into 19 groups labeled with different colors. The other DNA binding domain TF group is a collection of DNA-binding TFs that do not fit into any of the other major domain-based TF groups. (C). Heat map representing the averaged differential expression level (AM/CL, log2 fold change) of 65 TFs across seven time points from three independent biological replicates data. These data were normalized by genes in each column. Genes labeled in green color represent the TF which have already been detected in signaling pathways. (D). Phenotypes and relative regenerative length of regenerated legs when nine regeneration related TFs were knocked down separately. The significance of differences was analyzed by two-tailed Student’s *t test*. *: *P* <0.05, **: *P* <0.01, ***: *P* <0.001. n=3.

We found that RNAi knockdown of *Dll* which encodes a homeodomain protein, and *Zn finger homeodomain 2* (*zfh-2*) which encodes a C2H2 zinc finger homeobox protein, disrupted the entire regeneration process. No femur or distal parts could be regenerated after RNAi knockdown each of these two genes. In addition, the *dac* and C2H2 zinc finger TF *<u>b</u>rother of <u>o</u>dd <u>w</u>ith entrails limited* (*bowl*) were demonstrated to have a strong influence on the patterning of the regenerated femur-tibia-tarsus, the joint sites among the femur, tibia and tarsus exhibited a swelling phenotype, whose phenomenon is similar to Notch signaling. Additionally, *Ets at 21C* (*Ets21C*), *LIM homeobox 1* (*Lim1*), *Ultrabithorax* (*Ubx*), *BarH2* (*B-H2*), and *<u>b</u>ric <u>a</u> <u>b</u>rac <u>1</u>* (*bab1*) are also important for the regeneration ability, especially the size of the regenerated legs. Finally, the RNAi screen has revealed that nine TFs play significant roles during leg regeneration (Fig. 3D, Fig. S4A and S4B). Together with the previous six positive TFs (*Stat*, *Grh*, *Med*, *Mad*, *Sd*, and *Ci*), approximately one quarter (15/65) of the screened TFs were confirmed to contribute to leg regeneration. Among the top four candidates, *Dll* has been reported as an indispensable TF for the formation of the tarsus, and *dac* is necessary for elongation of the tibia and formation of the most proximal tarsomere in the cricket *G. bimaculatus* ^15,19^, while the phenotypic defects of *Dll* and *dac* in cockroach leg regeneration are much stronger than those in crickets. We then focused on the remaining two less well-known regeneration-associated TF genes, *zfh-2* and *bowl*, in the following research.

### *zfh-2* Contributes to Blastema Cell Proliferation under the Control of BMP and JAK-STAT Signaling

*zfh-2*, as a TF with almost the strongest phenotype in our RNAi screening results, was selected for in-depth study. RNAi knockdown of *zfh-2* treatments were performed at multiple time points (0, 3, and 5 dpa), and gradually decreased regenerative defects were observed when knockdown at later time points. When dsRNA injection and amputation were performed simultaneously (0 dpa), basically no visible new leg could be regenerated (Fig. 4A). While when RNAi was performed at 3 dpa, only two tarsus segments with defective in joint formation were regenerated, there was no obvious defect on femur or tibia. This strong influence of the *zfh-2* was confirmed using a second dsRNA (ds*zfh-2* 2nd) against different regions of this gene (Fig. S5A). To validate the notable impact of *zfh-2* through RNAi at 0 dpa, μ-CT scanning was performed, revealing no obvious new regenerated leg at 10 dpa in ds*zfh-2* group (Fig. S5B). This timeframe represents a late stage, typically characterized by the clear emergence of a well-defined new leg shape (Fig. 1B). Meanwhile, RNAi knockdown of *zfh-2* significantly reduced the number of mitotically active cells in the blastema when measured with EdU and PH3 staining (Fig. 4B and S5C). These results suggest that *zfh-2* controls blastema cell proliferation at early stage.

**Fig. 4.**
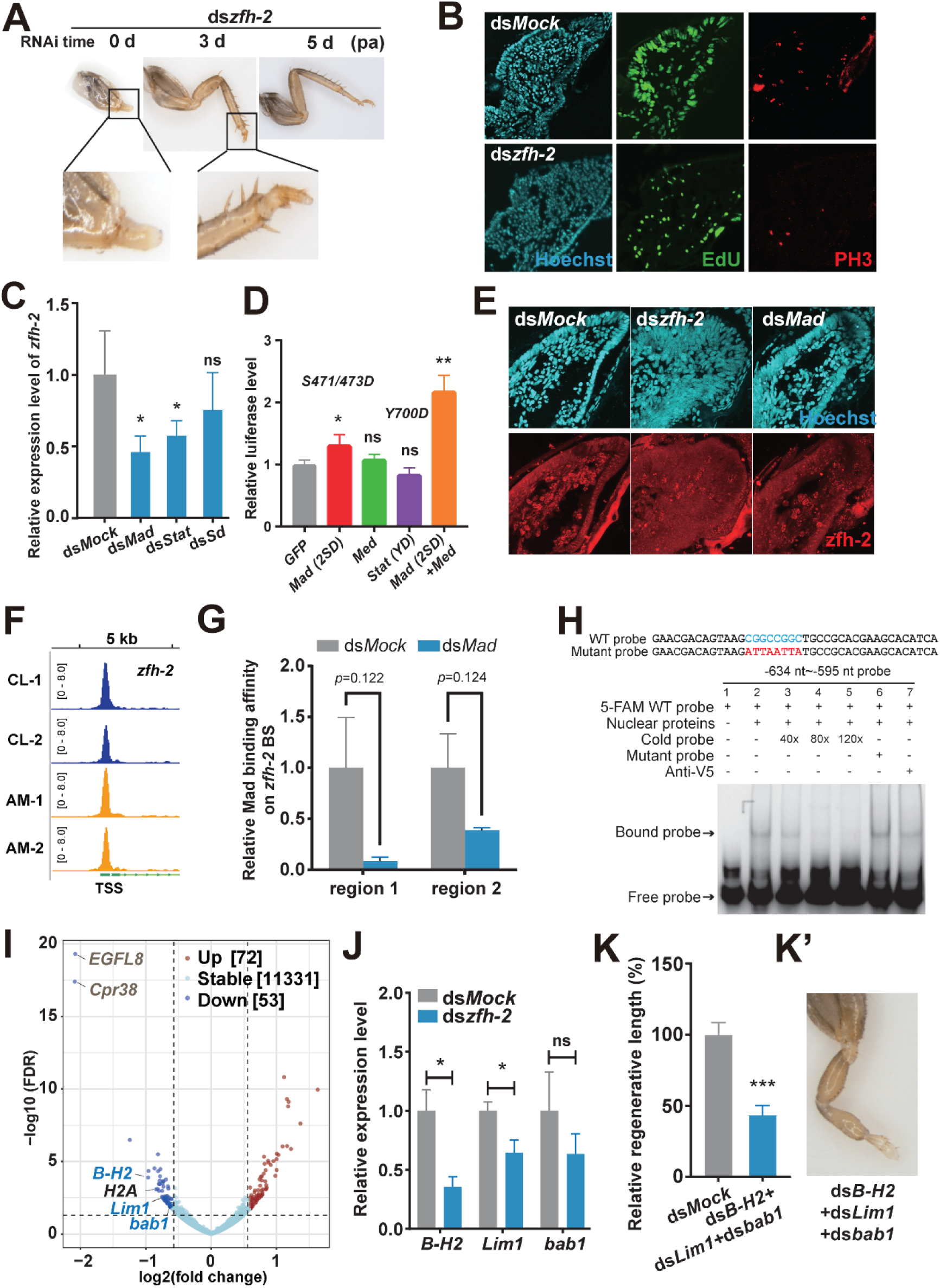
*zfh-2* contributes to blastema proliferation and morphogenesis under the control of the BMP and JAK-STAT signaling. (A). Phenotypes of regenerated legs after ds*zfh-2* treatments at three different time points. The legs in bottom line are zoom magnified from up line. n=3. (B). EdU and PH3 staining of proliferation positive cells. dsRNAs were injected at 0 dpa and the samples were harvested at 3 dpa, the EdU was injected at 6 h before harvest. n=3. (C). qRT‒PCR detection of the relative expression of *zfh-2* under the dsRNA treatments of *Mad*, *Stat* and *Sd*. n=3. (D). Utilizing a dual luciferase assay to detect the regulatory impact of proteins on the *zfh-2* promoter. n=3. (E). Immunohistochemistry staining of pMad and zfh-2 protein. (F). IGV snapshot of *zfh-2* displaying chromatin accessibility on its TSS region during unamputated (CL) and amputated (AM) conditions. (G). CUT&Tag qPCR detection of the relative Mad protein binding affinity on zfh-2 BS in both ds*Mock* and ds*Mad* groups, two replicates were used. Data are mean±sd, the differences were analyzed by two-tailed Student’s *t*-test. (H). EMSA using overexpressed Mad(2SD)/Med proteins from KC cells incubated with FAM-labelled WT probe and additional unlabeled probes (cold and mutant). All the probes containing a putative binding site (in blue) were derived from the region −634—595 nt distance to ATG, and the mutant nucleotides are marked in red. (I). Volcano plot analysis of the DEGs under ds*zfh-2* treatment. The violet dots indicate downregulated genes, and the red dots indicate upregulated genes. (J). Relative down regulation of *B-H2*, *Lim1*, and *bab1* under ds*zfh-2* treatment. (K-K’). Phenotypes and relative regenerative length of regenerated legs when *B-H2*, *Lim1*, and *bab1* were knocked down simultaneously. n=3. The significance of differences was analyzed by two-tailed Student’s *t test*. *: *P* <0.05, **: *P* <0.01, ***: *P* <0.001, “ns” stands for no significant difference.

To further explore the latent interaction of major signaling pathways with *zfh-2* during blastema cell proliferation, the BMP, JAK-STAT, and Hippo pathways were selected for examinations as their phenotypes are similar to *zfh-2*, and BMP and JAK-STAT control cell proliferation as well. Interestingly, the expression of *zfh-2* was reduced significantly when the *Mad* and *Stat* were silenced, while no significant change was observed when *Sd* was silenced (Fig. 4C). These results suggest that *zfh-2* controls cell proliferation as a downstream effector of the BMP and JAK-STAT signaling in the blastema. To further detect whether the Mad or Stat protein directly regulates the *zfh-2* promoter, the phosphomimic proteins Mad(2SD) and/or Stat(YD) were overexpressed in *Drosophila* Kc cells and dual luciferase reporter assay was performed. There was no significant change in promoter activity when Stat(YD) was overexpressed, which indicates that *zfh-2* is likely not the direct target of phosphorylated Stat. The Mad(2SD) alone upregulated the activity of *zfh-2* promoter, and co-expression of the Med, also named as co-Smad, enhanced the activity of Mad(2SD) ^28^, but Med alone had no detectable effect (Fig. 4D). Besides, phosphorylated Mad (pMad) and zfh-2 proteins with similar wild spread distributions in regenerating cells detected by serial section staining (Fig. S4D), and the protein level of zfh-2 decreased under ds*zfh-2* and ds*Mad* treatments (Fig. 4E and S5E). Next, the chromatin accessibility around transcription starting site (TSS) site was detected using ATAC-Seq, and obvious peaks were observed under both unamputated CL group and amputated AM group (Fig. 4F). This observation suggests that the chromatin is open in both conditions, the regulation of gene expression may depend on the presence of specific regulators, such as pMad protein. Furthermore, a potential Mad binding site (BS) was found inside of the peak by using JASPAR database (https://jaspar.elixir.no/). The binding affinity between pMad protein and *zfh-2* promoter BS in regenerating tissues was detected by using CUT&Tag qPCR, the Mad protein binding affinity on two different regions around the BS decreased in ds*Mad* group compared with ds*Mock* group (*p*=0.122 and *p*=0.124, respectively) (Fig. 4G). Dual luciferase reporter assay was used again to confirm the existence of binding site, no obvious activity was observed when the binding site was deleted (Fig. S5F). Moreover, the electrophoretic mobility shift assay (EMSA) was performed to detect the direct protein binding *in vitro*, and a specific binding band which can be competed by cold probe but not mutant probe was observed (Fig. 4H). These results together indicate that *zfh-2* might be the direct target of phosphorylated Mad protein during leg regeneration in cockroach.

Another RNA-seq was performed to search for the genes downstream of *zfh-2* (Table. S2). Fifty-three downregulated genes, including three TF genes, *B-H2*, *Lim1*, and *bab1*, were found after *zfh-2* RNAi treatment (Fig. 4I). It is noteworthy that these three TFs have been demonstrated to contribute to leg regeneration (Fig. 3D and S4B). The chromatin accessibility around the TSS regions of these three TF genes were also analyzed using the ATAC-Seq data. Strong chromatin accessibility peaks were observed in both unamputated CL group and amputated AM group of *B-H2*, while the accessibility of *Lim1* and *bab1* were significantly enhanced in the amputated AM group compared to the unamputated CL group (Fig. S5G). These peak regions may contain the cis-elements of their upstream regulators. When these three genes were simultaneously knocked down (Fig. S5H), a significantly more pronounced regeneration defect occurred, the overall size was much small and the segment number of tarsus was reduced (Fig. 4K-4K), surpassing the severity observed when RNAi was applied to each gene separately (Fig. 3D). In addition, the eight top downregulated non-TF genes (*Svil*, *Cpr38*, *EGFL8*, *Grcw*, *GDF10*, *Grcw2* and *H2A*) were also silenced *in vivo*, and no significant phenotypic defects were observed after RNAi knockdown of any factor, except for *H2A*, which died during molting with no regenerated leg visible (Fig. S5I). In brief, *zfh-2* is a crucial TF gene that transduces upstream BMP and JAK-STAT signaling to three downstream TFs via the BMP/JAK-STAT-*zfh-2*-*bab1*/*B-H2/Lim1* transcriptional cascade that contributes to leg regeneration.

### *drm* and *bowl* Regulate Leg Morphogenesis under the Control of Notch Signaling

The *bowl* was also selected due to its significant effect on morphogenesis and patterning (Fig. 3D). To confirm its function in leg regeneration, a second dsRNA against a different region (ds*bowl*-2) was designed and injected. An almost identical phenotype was observed, characterized by a significantly smaller leg and a failure of tissue morphogenesis and segment joint formation, and an ectopic outgrowth at the femur-tibia joint was observed (Fig. S6A, S6A’, and Fig. 5A’, arrow). To identify the regeneration stage at which *bowl* might function, a series of RNAi treatments were performed at multiple time points (0, 1, 2, 3, 5, 7 dpa). The strong phenotype was observed when the dsRNA injections were performed at 0 dpa and 1 dpa; thereafter, the phenotypes gradually weakened until 5 dpa (Fig. 5A), suggesting that *bowl* might function during the early-middle regenerative stages. The decreased level of bowl protein resulted in a much smaller new regenerated leg inside the exoskeleton in the ds*bowl* group compared with that in the ds*Mock* group detected by whole mount tissue staining and μ-CT scanning (Fig. 5B-5C and S6B). EdU and PH3 labeling indicated a slight decrease in cell proliferation in the blastema of the *bowl* knockdown animals (Fig. S6C and S6C’), explaining the smaller size of the regenerated legs. To elucidate how *bowl* affects leg patterning, RNA-seq was conducted after *bowl* RNAi, revealing 165 upregulated and 31 downregulated DEGs (Fig. 5D, Table. S3). Biological process GO enrichment was performed for the downregulated DEGs, revealing various developmental and morphogenesis processes, such as multicellular organism development, anatomical structure morphogenesis, and system development processes (Fig. 5E, Table. S4). Interestingly, *bab1*, previously verified as a TF influencing leg regeneration (Fig. 3D), was identified as a downstream effector of *bowl* (Fig. 5D and 5F), indicating a regulatory relationship between these two TFs during leg regeneration.

**Fig. 5.**
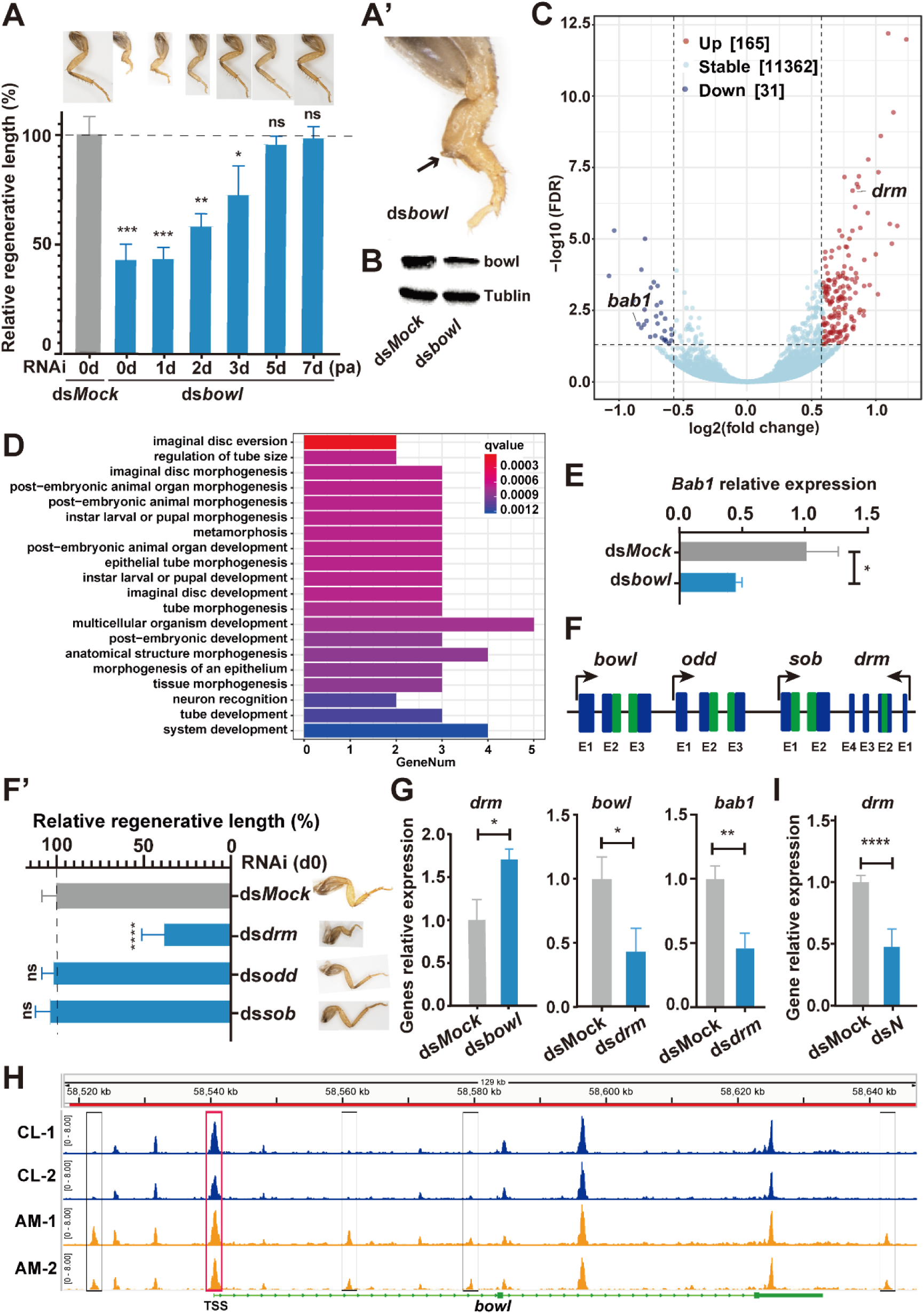
The odd-skipped family members *bowl* and *drm* regulate leg morphogenesis in a feedback loop under the control of Notch signaling. (A). Phenotypes and relative regenerative length of regenerated legs when *bowl* was knocked down at six different time points post amputation. n=3. (A’). The magnified picture of regenerated leg after ds*bowl* treatment at 0 dpa. The arrow indicates the ectopic outgrowth at the femur-tibia joint. (B). Protein level detection of *bowl* under RNAi using Western blotting. (C). Volcano plot analysis of the DEGs under ds*bowl* treatment condition. The violet dots indicate downregulated genes, and the red dots indicate upregulated genes. The ds*bowl* was injected at 0 dpa and performed for RNAi-Seq 3 days post injection. (D). Barplot of biological processes after GO enrich of downregulated DEGs. (E). Relative expression of *bab1* after ds*bowl* treatment which measured by FPKM values. (F-F’). Mapping of odd-skipped family member genes *bowl*, *odd*, *sob*, and *drm* on the chrome and the phenotypes after dsRNA treatments. Exons are represented by dark blue rectangles, green rectangles stand for the conserved region among these gene, black arrows represent the direction of gene transcription (F); phenotypes and relative regenerative length of regenerated legs after ds*drm*, ds*odd*, and ds*sob* treatments at d0 (F’). (G). Relative expression of genes under ds*bowl* or ds*drm* treatments detected by qRT‒PCR. n=3. (H). IGV snapshot of *bowl* displaying accessibility on its TSS and other regions during developmental (CL) and regenerative (AM) stages. (I). Relative expression of *drm* under ds*N* treatment detected by qRT‒PCR. n=3. The significance of differences was analyzed by two-tailed Student’s *t test*. *: *P* <0.05, **: *P* <0.01, ***: *P* <0.001, “ns” stands for “no significant difference”.

The *drumstick* (*drm*), also a member of the odd-skipped family, was identified with a high fold change among the upregulated DEGs (Fig. 5D). Together with *<u>s</u>ister of <u>o</u>dd and <u>b</u>owl* (*sob*) and *odd-skipped* (*odd*) ^29^, all four members of the odd-skipped family are clustered on the same chromosome. Surprisingly, these four genes exhibit highly conserved DNA and amino acid sequences within their DBDs (Fig. 5G and S6D). Therefore, we separately knocked down *drm*, *odd*, and *sob* to explore their potential functions in leg regeneration, and only *drm* RNAi significantly influenced the leg size and patterning (Fig. 5G’). The phenotypic defects observed after *drm* RNAi were similar to those observed after *bowl* RNAi, and a similar ectopic outgrowth at the femur-tibia joint was observed as well (Fig. S6E and S6E’, arrow). These results imply that *bowl* and *drm* might function in regeneration through an interrelated mechanism. The upregulation of *drm* after *bowl* RNAi was confirmed by qRT-PCR, whereas the expression levels of *bowl* and *bab1* significantly decreased under *drm* RNAi treatment (Fig. 5H). The composite experimental data suggest that *bowl* functions as a repressor of *drm* expression, and *drm*, in turn, acts upstream of *bowl* and *bab1* as an activator, forming a transcriptional cascade. The chromatin accessibility around the TSS region of the *bowl* was also detected using ATAC-Seq, revealing strong accessibility in both unamputated and amputated conditions. Moreover, in the amputated group, two peaks were gained on the outer sides of gene region, and two peaks were gained within the introns of the *bowl* (Fig. 5I), suggesting increased accessibility of certain enhancers during regeneration. In addition, we searched for candidate signaling pathways upstream of *drm* and identified Notch signaling due to the similar abnormal morphogenetic phenotypes. Intriguingly, the expression level of *drm* decreased significantly after *N* gene RNAi treatment, showing that *drm* functions downstream of Notch signaling during morphogenesis (Fig. 5J). In short, the *drm* and *bowl* work in a feedback loop, and a Notch-*drm*/*bowl*-*bab1* cascade contributes to morphogenesis during leg regeneration.

In summary, our research reveals that *zfh-2-* and *bowl* are two crucial TFs for cockroach leg regeneration. Moreover, we have discovered that signaling pathways and TFs interact with each other, forming a regulatory network that contributes to distinct processes during leg regeneration. Furthermore, our findings demonstrate that two step-by-step transcriptional cascades, BMP/JAK-STAT-*zfh-2*-*bab1*/*B-H2/Lim1* and Notch-*drm/bowl*-*bab1*, play predominate roles in blastema cell proliferation and morphogenesis.

## Discussion

Here, we chose the American cockroach as our research object. The regeneration phenomenon of this species was initially documented in the 1840s ^30^, and it has since become a prominent insect model in the field of regenerative biology. Cockroaches exhibit a top-ranked regeneration ability among Insecta; in addition to the leg, all appendages of the cockroach can be regenerated after amputation. Moreover, the classical leg transplantation experiment was initially performed on cockroaches ^31^. Research on cockroach regeneration had recently come to a standstill due to the absence of reference genomic data, a challenge that has been addressed by our team and collaborators ^14^. Now, the American cockroach can be served as an ideal insect model for appendage regeneration. It boasts several advantages, including ease of raising, distinct leg shape compared to *Drosophila* imaginal discs, a short regenerative period, sensitivity to dsRNA, flexibility for *in vivo* screening (overcoming potential gene knockout-induced embryonic death) ^32^, and a feasible gene knockout strategy ^33^. In this study, firstly, leveraging advanced μ-CT scanning and tissue staining technologies, we provided the most intensive and detailed morphological profiles of internal microstructures and cellular patterning during insect appendage regeneration to date. Secondly, many signaling pathways (including JAK-STAT, BMP, Wnt/Wg, Hh, Hippo, EGFR) and TFs (such as *Dll* and *dac*) that have been demonstrated to be necessary for leg regeneration in American cockroaches have also been found to play important roles in tissue regeneration, such as tail regeneration in tadpoles ^34–36^, gill and limb regeneration in newts and axolotls ^37,38^, and liver regeneration in mice ^39^. These results indicate that the roles of signaling pathways and TFs in appendage regeneration are evolutionarily conserved from cockroaches to vertebrates. In addition, an ancient and common satellite cell-driven mechanism for muscle regeneration in arthropods and vertebrates has been reported ^40^, further supporting the use of the cockroach as an ideal animal model for regeneration research from molecular and evolutionary perspectives. Within complex regeneration processes, many signaling pathways and TFs are needed to be precisely regulated to ensure that they are expressed in the desired cells, at the right times, and in the right amounts. Transcriptional regulation is a critical mechanism that allows organisms to respond to the signals induced by appendage loss. The transcriptional cascade BMP/JAK-STAT-*zfh-2*-*bab1*/*B-H2/Lim1*, was found to control both blastema cell proliferation and morphogenesis. In this cascade, the C2H2 zinc finger protein *zfh-2* plays a pivotal role in transducing the upper JAK-STAT and/or BMP signals to downstream TFs. In this study, *zfh-2* was identified as functioning in both blastema cell proliferation and tarsus segmentation during leg regeneration for the first time (Fig. 4). In *Drosophila*, *zfh-2* is required for proximo-distal patterning and tarsus segmentation as a downstream factor of Notch signaling instead of Dpp signaling during leg development ^41^. We hypothesize that *zfh-2* may also be regulated by Notch signaling in the late stage of cockroach leg regenerative morphogenesis. This speculation arises from our observation that *zfh-2* is required for tarsus regeneration at a relatively late stage (Fig. 4A). The potential disparity in the regulation of *zfh-2* by the Dpp signal between *Drosophila* leg development and cockroach leg regeneration could be elucidated through an evolutionary developmental perspective in future. Moreover, this gene is crucial for wing hinge development and is essential for ionizing radiation-induced regenerative behavior in wing discs. It functions as a downstream effector of the JAK-STAT pathway to promote the survival of JNK-signaling cells ^42,43^. In addition, as a continuation of our previous work showing the essential role of *Dpp* and *Mad* in cockroach leg regeneration ^14^, *zfh-2* was revealed to be a target of the phosphorylated Mad protein during blastema cell proliferation to a certain degree. Since *zfh-2* can transduce both the upstream JAK-STAT and/or BMP signals, further studies are required to clarify the relationship between these two pathways in the context of regeneration in the future. As important down regulators of *zfh-2*, the TFs *B-H2*, *bab1*, and *Lim1* were also newly discovered to control leg regeneration. Moreover, *zfh-2* represents a potentially influential gene associated with appendage regeneration in other animals: the vertebrate ortholog *Zfhx4* is upregulated by about 1.8-fold in the caudal fin regeneration of the zebrafish, *Danio rerio* ^44^, about 1.6-fold in the tail regeneration of the tadpole, *Xenopus tropicalis* ^45^, and approximately 4.0-fold in the limb regeneration of the newts, *Plethodontid salamander* ^46^ (Fig. S7A). However, the roles of vertebrate orthologs of *zfh-2* in appendage regeneration have not been examined, and further attempts are required to validate their potentially conserved regenerative function.

In addition to the six signaling pathways discussed above, we here discovered that Notch signaling promotes morphological changes associated with leg size and joint formation during leg regeneration via the Notch-*drm/bowl*-*bab1* transcriptional cascade (Fig. 5). Notch signaling has been reported to play a role in fin and heart regeneration in zebrafish ^47,48^, tail regeneration in tadpoles ^49^, and retinal regeneration in newts ^50^. In insects, Notch signaling was observed to be activated during leg regeneration in *Harmonia axyridis* ^13^ and involved in cell proliferation during *Drosophila* wing discs and intestinal regeneration ^51,52^. Notably, this represents the first report of Notch signaling directly controlling leg regeneration in insects. Interestingly, we have observed that the odd-skipped family TFs *drm* and *bowl* function downstream of Notch signaling in a feedback loop to promote morphological patterning. In vertebrates, the *<u>o</u>dd <u>s</u>kipped-<u>r</u>elated* genes (*Osr1*/*Osr2*), the ortholog of *bowl*, are upregulated about 2.8-fold in tadpole tail regeneration ^45^ and about 2.7-fold in newt limb regeneration ^46^ (Fig. S7B). Besides, the *Osr1* is utilized as a marker for fibro-adipogenic progenitors during myogenesis and muscle regeneration in mammals ^53,54^. This information suggests that the orthologs of *bowl* might also play a crucial role in appendage regeneration in vertebrates. The relationship between Notch signaling and odd-skipped genes during developmental leg segmentation has been reported in *Drosophila* ^29,55^, and the TF *Bab* has also been found to be required for developmental specification and proper segmentation of the tarsus in *Drosophila* ^56^. In addition, *bab1* has been newly identified as essential for leg regeneration downstream of *bowl* in this cascade (Fig.5F), and further investigation is needed to unravel its detailed transcriptional regulatory mechanism. Interestingly, *bab1* is involved in both the *bowl-* and *zfh-2-*driven transcriptional cascades, forming a regulatory network that underscores the complexity and intricacy of leg regeneration regulation.

### Limitations of the study

Our study primarily focused on the transcriptional regulation of leg regeneration. Consequently, certain aspects were inevitably overlooked, and these can be addressed in future studies. Firstly, to clarify the main focus of this work, we paid more attention to the signaling pathways, and the large-scale screening of essential regeneration genes also focused on TFs. Thus, the analysis and mining of RNA-Seq data were insufficient. For instance, we paid almost all our attention to TFs and associated pathways, other gene categories outside the scope of this study (e.g. catalyzing enzymes and kinases) are also worth exploring. Secondly, due to technical limitations in this non-model species, *P. americana*, no gene activation or overexpression strategy can be utilized at the moment. Thus, our results indicated that these two transcriptional cascades are necessary for leg regeneration, while it remains challenging to understand how these cascades control the regenerative processes. Thirdly, our results showed that these two transcriptional cascades predominantly orchestrate blastema cell proliferation and/or morphogenesis, but it is not clear which kinds of cell types are involved in these processes. The use of more in-depth and high-resolution technologies, such as single-cell/nucleus sequencing and spatial transcriptomics, will significantly aid in addressing this question. In addition, the insect regeneration might be distinct from vertebrate models at a certain degree, such as the insect blastema is derived from epidermal cells whereas the vertebrate blastema is from mesenchyma ^59^. Our newly discovered transcriptional cascades, BMP/JAK-STAT-*zfh-2*-*bab1*/*B-H2/Lim1* and Notch-*drm/bowl*-*bab1*, have not yet been identified in vertebrate appendage regeneration, it will be interesting to investigate whether these two cascades evolutionarily conserved regulate appendage regeneration in vertebrate animal systems. Moreover, even though the models of both tissue and gene expression patterning have been reported to diverge between regeneration and development in arthropods ^51,58^, many of the signaling pathways and key genes essential for development might also be essential for appendage regeneration. This information indicates the complex relationship in the regulatory mechanisms between appendage development and regeneration, and further attempts are required to clarify their similarities and differences in the future.

Overall, we show that two transcriptional cascades orchestrate leg regeneration in the American cockroach, an ideal insect model for appendage regeneration research. The progress made with this model will provide a broader context for understanding regeneration mechanisms from an evolutionary perspective.

## Methods

### Key resources table

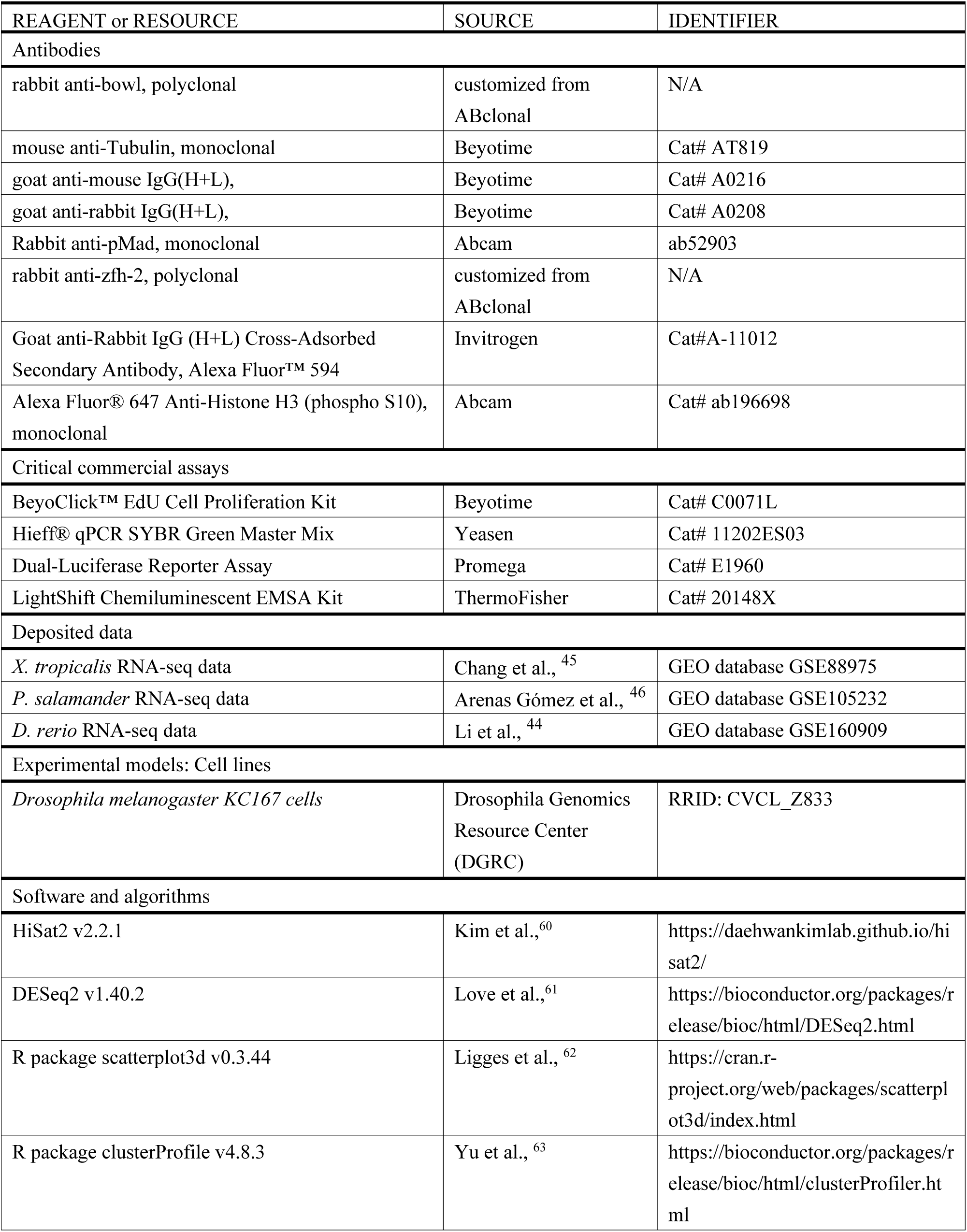
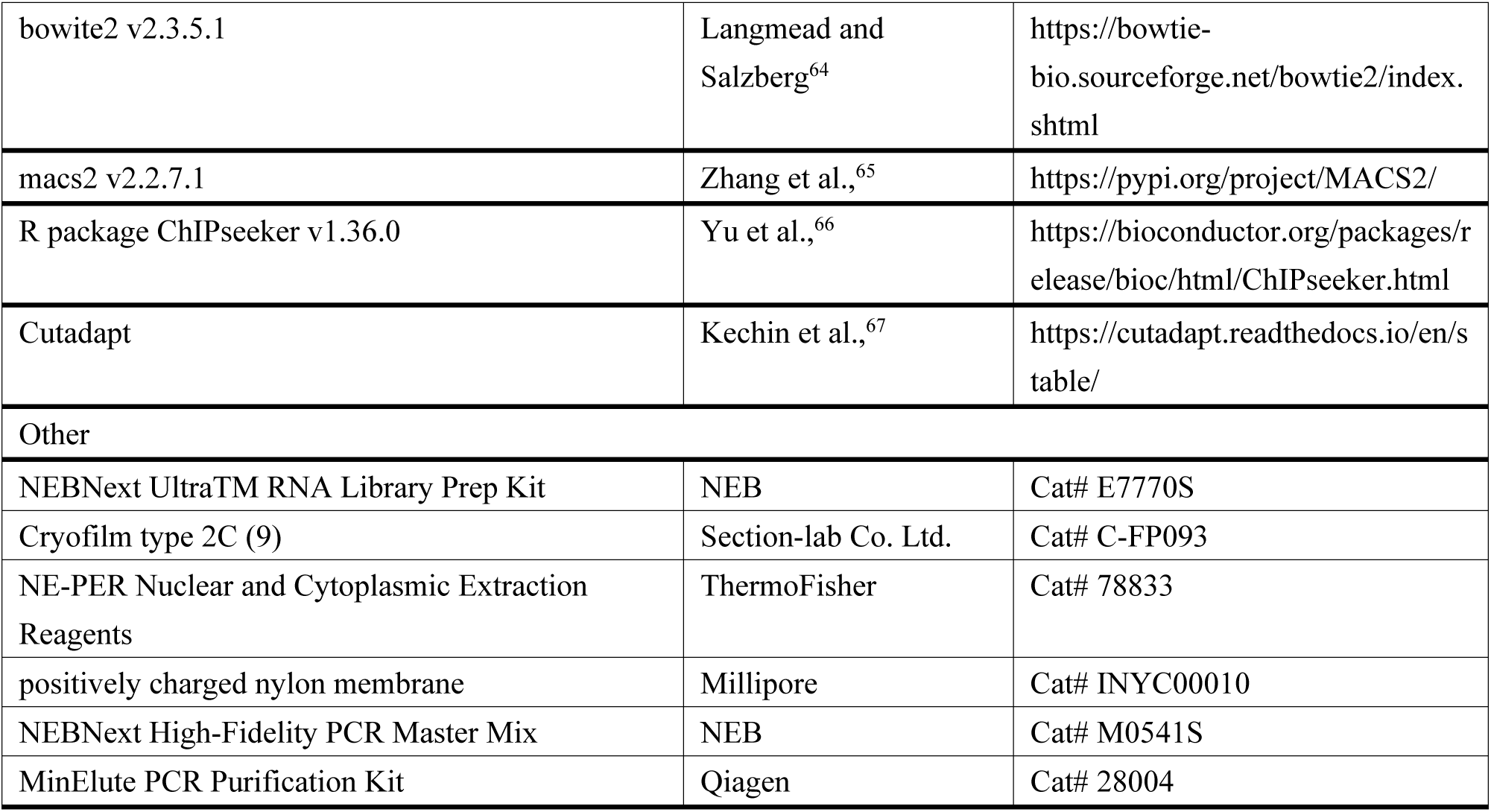

### Resource availability

#### Lead contact

Further information and requests for resources and reagents should be directed to and will be fulfilled by the lead contact, Sheng Li.

#### Materials availability

The materials are available by connecting the lead contact.

#### Data and code availability

All RNA-Seq and ATAC-Seq data are available via the Sequence Read Archive (SRA) database under accession numbers PRJNA868495.

This paper does not report original code. We used publicly available software in this study. Please see the METHOD DETAILS section for further details.

Any additional information required to reanalyze the data reported in this work paper is available from the lead contact upon request

### Experimental model and study participant details

The line of *P. americana* was provided by Dr. Huiling Hao, and this line has been maintained through inbreeding for 30 years. The colony was housed at 27 °C with a relative humidity of 70–80% in plastic cages, and the animals were provided with commercial rat food and water ad libitum. To obtain pools of synchronized animals, newly molted 4^th^ instar nymphs were selected from the colony and placed in separate containers, where they were supplied with water and rat food.

### Method details

#### Leg amputation and regenerative length calculation

Amputation operations were performed on CO_2_-anaesthetized 4^th^-6^th^ instar animals at 1-day post molting. The distal tarsus, tibia and femur of one metathoracic leg (hindleg) was removed at the junction of the trochanter and femur, and trochanter and coxa were left to observe regeneration. The contralateral hindlegs served as internal controls for analyzing the regenerative length. After another molting, the length of regenerated and contralateral tarsus, tibia and femur were measured. The relative regenerative lengths were calculated by dividing the length of the regenerated legs by the length of the contralateral legs.

#### CT scanning and 3D model construction

All regenerating legs used for X-ray μ-CT were fixed with neutral paraformaldehyde solution and dehydrated in pure n-propanol, then in ethanol solutions from 75 to 100%, stepwise. They were critical-point dried (Leica EM CPD 300). The specimens were scanned by an X-radia Micro CT-400 scanner at the Institute of Zoology and at Xishuangbanna Tropical Botanical Garden, Chinese Academy of Sciences (beam strength: 60KV, absorption contrast). The resolution is about 5 μm. The three-dimensional (3D) reconstruction model of regenerating legs were reconstructed and smoothed with Amira 5.4 based on the obtained image stacks from μ-CT. Final figures were prepared with Adobe Photoshop (CS6).

#### Paraffin sections and histological HE staining

The samples were harvested at 6 h, 12 h, 1 d, 3 d, 5 d, 7 d and 10 d post amputation and subsequently fixed with paraformaldehyde solution for 48 h. Thereafter, the samples were washed with phosphate-buffered saline for 30 min three times and dehydrated via an alcohol gradient with incubation for 30 min at each step. Dimethylbenzene was used to clear the samples for 30 min. Finally, the samples were incubated in wax for an additional 6 h. Paraffin sections (approximately 7 μm) were performed to make the sections containing the coxa and trochanter as complete as we could. Staining was performed with hematoxylin and eosin (H&E). The slides were evaluated by light microscopy (Evos, FL Auto, Invitrogen).

#### Cell proliferation assay

Cells proliferation were detected using the BeyoClick™ EdU Cell Proliferation Kit with Alexa Fluor 488 (Beyotime, C0071L) and anti-Histone H3 (phospho S10) antibody [Abcam, ab196698]. 1 μL of 5 mM EdU was injected into the abdomen of 4^th^ instar nymphs 6 hours before sample harvest, and the samples were subsequently fixed with paraformaldehyde solution for 48 h. Thereafter, the samples were washed with phosphate-buffered saline for 30 min three times and dehydrated via 50 % sucrose with incubation for 24 h. The frozen section was performed with the assistance of Cryofilm type 2C (9) (Section-lab Co. Ltd., C-FP093) to prevent the sample with crunchy cuticles from elution from the slide. The sections were incubated with PBT (PBS with 0.3% TritonX-100 and 0.5% BSA) for 2 h. Blocked samples were incubated with primary antibody Alexa Fluor® 647 Anti-Histone H3 (phospho S10) antibody at 1:300 in PBT for 1 h at room temperature. The samples were washed with PBT for three times. EdU-incorporated cells were detected according to the manufacturer’s instructions. Finally, the samples were washed with PBT and incubated with a 1:1000 dilution of Hoechst 33342 in PBT for 15 min for nuclear staining. For ds*bowl* and ds*zfh-2* treatments, the dsRNA were injected at 0 dpa and the tissues were harvested at 3 dpa.

#### RNA-Seq and data analysis

For differential expressed gene (DEG) pattern detection during regeneration, the first amputations were performed on one of the hindlegs at the first day post molting, after that, the remaining distal half coxa and the whole trochanter at 6 h, 12 h, 1 d, 3 d, 5 d, 7 d, and 10 d post amputation in both experimental (AM) and contralateral (CL) sides were harvested. The samples from about 20 individuals were dissected and mixed to form one sample, and three replicates were performed for each treatment. For differential expressed gene detection after RNAi of target genes, the dsRNA injection was performed together with amputation, and the samples were harvested at 3 dpa. A total of 1 μg of RNA per sample was used as input material for RNA preparation. Sequencing libraries were generated using a NEBNext Ultra^TM^ RNA Library Prep Kit for Illumina (NEB, E7770S) following the manufacturer’s recommendations. Clustered libraries were sequenced on an Illumina platform and paired-end reads were generated. Raw FASTQ-format data were first processed through in-house Perl scripts. HiSat2 v2.2.1 software was used to map the reads to the reference genome (pame.OGS1.v2) ^60^. Gene expression levels were estimated as fragments per kilobase of transcript per million fragments mapped (FPKM) values, and differential expression analysis between the AM and WT groups was performed using DESeq2 v1.40.2 ^61^. Genes with adjusted *P*-values < 0.05 in DESeq2 were selected. Principal component analysis (PCA) was plotted using the R package scatterplot3d v0.3.44 ^62^. Gene Ontology (GO) and Kyoto Encyclopedia of Genes and Genomes (KEGG) pathway analysis were performed using the R package clusterProfiler v4.8.3 ^63^ for the DEGs. Genes within certain clusters displayed similar expression profiles, barring minor variations.

#### Quantitative real-time PCR (qRT*‒*PCR) for gene expression detection

For gene expression analyses, qRT‒PCR was performed in triplicate using Hieff® qPCR SYBR Green Master Mix (Yeasen, 11202ES03). Relative gene expression was calculated using the ΔΔCt method following the manufacturer’s instructions. The primers used for qRT‒PCR are shown in Table S5. All the specificities of the primers were confirmed by Sanger sequencing of the PCR products.

#### Double-stranded RNA (dsRNA) treatment

For negative control, a ds*Mock* targeting a clone vector sequence was used ^14^. Sense and antisense RNA was synthesized in a single reaction using a T7 RiboMAX Express RNAi System (Promega, P1700). After purification, the dsRNA concentration was adjusted to 2 μg/μl. Under normal conditions, nymphs were injected with about 2 μg of dsRNA between the abdominal sternites using a syringe for each injection. The injection time points will be remarked in the figure legends. The simultaneous dsRNA knockdown of *B-H2*, *Lim1* and *Bab1* genes was implemented by injecting the mixture of three dsRNA solution, and 2 μg of each kind of dsRNA was injected into one animal. To eliminate potential off-target effects of our dsRNA, we replicated the experiment using a second dsRNA for dsz*fh-2* and ds*bowl*. The primer sequences for dsRNA synthesis can be found in Table S6.

#### Western blotting

For Western blotting analysis, the samples were dissected for the subsequent procedures. Proteins extracted with RIPA lysis buffer (Beyotime, P0013B) were run on polyacrylamide gels and transferred onto PVDF membranes for primary antibody incubation. The primary antibody anti-bowl (rabbit polyclonal antibody, 1:2000, customized from ABclonal) were used to detect the level of these two proteins, mouse anti-Tubulin (Beyotime, AT819) was used as a housekeeping protein antibody for normalization (1:5000). The secondary antibodies goat anti-mouse IgG(H+L) (1:5000) (Beyotime, A0216) and goat anti-rabbit IgG(H+L) (1:5000) (Beyotime, A0208) were used. The membranes were then treated with enhanced chemiluminescence reagent and the blots were captured by automatic chemiluminescence analysis system (Tanon, 5200 Muilti).

#### Immunohistochemistry

To detail the distribution of pMad and zfh-2 protein at the regenerating site, serial section staining was performed because both the primary anti-pMad antibody (Abcam, ab52903) and anti-zfh-2 antibody (rabbit polyclonal antibody, customized from ABclonal) were from rabbit. The primary antibodies were diluted 1:200 and the secondary anti-rabbit antibody (Alexa Fluor 594 or 488 coupled, Invitrogen) was diluted 1:400. The nuclei were stained with Hoechst 33342 (Beyotime, C1022).

#### Dual luciferase reporter assay

The promoter and 5’UTR of *zfh-2* was cloned into the pGL3-basic plasmid which containing the firefly luciferase gene between *Kpn*I and *Xma*I enzyme sites. We generated a Stat(YD) that mimics phosphorylation of Stat with Y700D mutant, and generated a Mad(2SD) that mimics phosphorylation of Mad with S471D and S473D mutants. The full length sequence of Med, Mad(2SD), Stat(YD), and negative control GFP genes were cloned into a p10xUAS plasmid which was promoted by actin-Gal4 plasmid. The pGL4 plasmid containing the renilla luciferase gene was used to normalize the transfection efficiency. The pGL3 plasmids with truncated prompter-UTR regions of *zfh-2* were cloned for the search of protein binding site. All these four plasmids actin-Gal4, p10xUAS, pGL3, and pGL4 were co-transfected into the *Drosophila melanogaster* KC167 cells in a 96-well plate. Three replicates were performed for each group. The activity of luciferases was measured by the kit of Dual-Luciferase Reporter Assay System (Promega, E1960).

#### ATAC-Seq and data analysis

Unamputated (CL) legs and amputated (AM) legs at 3 dpa were harvested. The samples from about 100 individuals were dissected and mixed to form one sample, and two replicates were performed for each treatment. In brief, native nuclei were purified from frozen samples. The Nextera DNA Library Preparation Kit (Illumina) was used to perform the transposition according to the manufacturer’s manual. 50,000 nuclei were pelleted and resuspended with transposase, for 30 min at 37 °C. The transposed DNA fragments were purified immediately after with a MinElute PCR Purification Kit (Qiagen, 28004). After samples were PCR-amplified using 1×NEBNext High-Fidelity PCR Master Mix (NEB, M0541S). Subsequent libraries were purified with the MinElute PCR Purification Kit (Qiagen, 28004) and subjected to sequencing on Illumina Novaseq 6,000 using PE150. The clean reads were trimmed with Cutadapt ^67^ and were then aligned to reference genome sequences using the bowite2 v2.3.5.1 program ^64^. The bam file generated by the unique mapped reads as an input file, using macs2 v2.2.7.1 software for callpeak with cutoff qvalue < 0.05 ^65^. Peaks were annotated by using R package ChIPseeker v1.36.0 ^66^. A custom Perl script was used to generate smoothened genome browser tracks in BigWig format for data visualization on the Integrative Genomics Viewer (IGV) v2.16.0 ^68^.

#### Electrophoretic mobility shift assay

The full length of Med and Mad(2SD) proteins were overexpressed together in KC cells, and the nuclear proteins were extracted using NE-PER Nuclear and Cytoplasmic Extraction Reagents (Thermo Fisher, 78833). The region −634–−595 nt fragment distance to initiation codon of the *zfh-2* was synthesized with FAM-labelling at the 5′ terminus, followed by denaturing and annealing to generate a labelled probe. Binding reaction was performed by incubating 8 μl of nuclear protein extracts with 100 μmol of FAM-labelled probe, in a total volume of 20 μl containing 1× binding buffer, 2.5% glycerol, 5 mM MgCl_2_, 0.05% NP-40 and 1 μg of poly(dI·dC). In the competition assays, 40-, 80-, and 120-fold molar excess of unlabelled (WT or mutant) probe was added into the binding reaction. For the super-shift assay, 1 μl of V5 antibody was preincubated with nuclear extracted proteins for 2 h at 4 °C before adding the labelled probe. All reactions were then maintained at room temperature for 20 min, and immediately separated by 6% native acrylamide gel electrophoresis. The gel was transferred onto a positively charged nylon membrane (Millipore, INYC00010), developed with LightShift Chemiluminescent EMSA Kit (Thermo Fisher, 20148X) according to the manufacturer’s instructions, and complexes were finally visualized via chemiluminescence.

#### CUT&Tag assay

The remaining distal half coxa and the whole trochanter injected with ds*Mock* or ds*Mad* at 3 dpa were harvested. CUT&Tag assay was performed using NovoNGS CUT&Tag 3.0 High-Sensitivity Kit (for Illumina, Novoprotein scientific Inc., Cat#N259-YH01). Briefly, ds*Mock* and ds*Mad* treated samples were fully lysed and counted up to 400,000 cells. Then 37% formaldehyde was gently added to cells (the final concentration of formaldehyde is 0.1%) and these samples were incubated at room temperature for 2 min. Cross-linking was stopped by addition of 2.5 M glycine for 5 min and then the samples were washed with the wash buffer. The cells were enriched by ConA Beads and resuspended by 50 µL primary antibody buffer of anti-pMad (1:50, Abcam, ab52903), and incubated overnight at 4 °C. We then discarded the primary antibody buffer and added 100 µL anti-rabbit IgG antibody buffer for 1 h at a dilution of 1:100. Then the beads were washed for 3 times using antibody buffer and then incubated with proteinA/G-Tn5 transposome for 1 h and washed for 3 times by ChiTaq buffer. Cells were resuspended in 50 µL Tagmentation buffer (10 mM MgCl_2_ in ChiTaq Buffer) and incubated at 37 °C for 1 h. The incubation was stopped by adding 10 µL 10% SDS at 55 °C for 2 h. The DNA fragments were extracted by Tagment DNA extract beads and amplified using 5× AmpliMix. Then the DNA was re-extracted by DNA clean beads for qPCR detection. The primers used for CUT&Tag qPCR are shown in Table S7.

### Quantification and statistical analysis

For relative regenerative length calculation, at least three animals were measured in each group, and the statistics were carried out on average values. Data are mean±sd, the differences were analyzed by two-tailed Student’s *t*-test. The data statistics in the paper were analyzed by software SPSS 25 and the specific analysis method was shown in the figure legends.

## Declarations

## Ethics approval and consent to participate

Not applicable.

## Consent for publication

Not applicable.

## Author contributions

C.R. and S.L. conceived the project. C.R. and S.L. designed and led the project. C.R., Y.W., S.Z. Z.Z., and E.Y.L. performed most of the experimental work and analyzed the data. C.Z. analyzed the μ-CT data. M. L., D. Y., and L. L. conducted bioinformatic data analysis. S. Liu, X. Z., W. W., and J. F. improved the manuscript. C.R., and S.L. wrote the manuscript.

## Acknowledgments

We would like to thank Ting Tang in Xishuangbanna Tropical Botanical Garden, Chinese Academy of Sciences, and Caixia Gao in Institute of Zoology, Chinese Academy of Sciences, for μ-CT scanning, thanks Mengting Huang in Taiyuan Normal University for the help of analysis of μ-CT data.

## References

1. Goss, R.J. (1969). Principles of Regeneration (Academic Press).

2. Nowoshilow, S., Schloissnig, S., Fei, J.F., Dahl, A., Pang, A.W.C., Pippel, M., Winkler, S., Hastie, A.R., Young, G., Roscito, J.G., et al. (2018). The axolotl genome and the evolution of key tissue formation regulators. Nature 554, 50–55. 10.1038/nature25458.

3. Maruzzo, D., Bonato, L., Brena, C., Fusco, G., and Minelli, A. (2005). Appendage loss and regeneration in arthropods: a comparative view. In Crustacea and Arthropod Relationships, Stefan, K., and Ronald, J., eds. (CRC Press), pp. 215–245. 10.1201/9781420037548.

4. Alwes, F., Enjolras, C., and Averof, M. (2016). Live imaging reveals the progenitors and cell dynamics of limb regeneration. Elife 5. 10.7554/eLife.19766.

5. Wang, J., Chen, X., Hou, X., Wang, J., Yue, W., Huang, S., Xu, G., Yan, J., Lu, G., Hofreiter, M., et al. (2022). “Omics” data unveil early molecular response underlying limb regeneration in the Chinese mitten crab, *Eriocheir sinensis*. Sci. Adv. 8, eabl4642. 10.1126/sciadv.abl4642.

6. Tanaka, E.M. (2016). The Molecular and Cellular Choreography of Appendage Regeneration. Cell 165, 1598–1608. 10.1016/j.cell.2016.05.038.

7. Zhong, J., Jing, A., Zheng, S., Li, S., Zhang, X., and Ren, C. (2023). Physiological and molecular mechanisms of insect appendage regeneration. Cell Regen. 12, 9. 10.1186/s13619-022-00156-1.

8. Galliot, B., Tanaka, E., and Simon, A. (2008). Regeneration and tissue repair: themes and variations. Cell Mol. Life Sci. 65, 3–7. 10.1007/s00018-007-7424-0.

9. Sood, P., Lin, A., Yan, C., McGillivary, R., Diaz, U., Makushok, T., Nadkarni, A.V., Tang, S.K.Y., and Marshall, W.F. (2022). Modular, cascade-like transcriptional program of regeneration in Stentor. Elife 11. 10.7554/eLife.80778.

10. Bosch, M., Serras, F., Martín-Blanco, E., and Baguñà, J. (2005). JNK signaling pathway required for wound healing in regenerating *Drosophila* wing imaginal discs. Dev. Biol. 280, 73–86. 10.1016/j.ydbio.2005.01.002.

11. Smith-Bolton, R.K., Worley, M.I., Kanda, H., and Hariharan, I.K. (2009). Regenerative growth in *Drosophila* imaginal discs is regulated by Wingless and Myc. Dev. Cell 16, 797–809. 10.1016/j.devcel.2009.04.015.

12. Mito, T., Inoue, Y., Kimura, S., Miyawaki, K., Niwa, N., Shinmyo, Y., Ohuchi, H., and Noji, S. (2002). Involvement of *hedgehog*, *wingless*, and *dpp* in the initiation of proximodistal axis formation during the regeneration of insect legs, a verification of the modified boundary model. Mech. Dev. 114, 27–35. 10.1016/s0925-4773(02)00052-7.

13. Zhou, H., Ma, Z., Wang, Z., Yan, S., Wang, D., and Shen, J. (2021). Hedgehog signaling regulates regenerative patterning and growth in *Harmonia axyridis* leg. Cell Mol. Life Sci. 78, 2185–2197. 10.1007/s00018-020-03631-7.

14. Li, S., Zhu, S., Jia, Q., Yuan, D., Ren, C., Li, K., Liu, S., Cui, Y., Zhao, H., Cao, Y., et al. (2018). The genomic and functional landscapes of developmental plasticity in the American cockroach. Nat. Commun. 9, 1008. 10.1038/s41467-018-03281-1.

15. Nakamura, T., Mito, T., Miyawaki, K., Ohuchi, H., and Noji, S. (2008). EGFR signaling is required for re-establishing the proximodistal axis during distal leg regeneration in the cricket *Gryllus bimaculatus* nymph. Dev. Biol. 319, 46–55. 10.1016/j.ydbio.2008.04.002.

16. Bando, T., Ishimaru, Y., Kida, T., Hamada, Y., Matsuoka, Y., Nakamura, T., Ohuchi, H., Noji, S., and Mito, T. (2013). Analysis of RNA-Seq data reveals involvement of JAK/STAT signalling during leg regeneration in the cricket *Gryllus bimaculatus*. Development 140, 959–964. 10.1242/dev.084590.

17. Herrera, S.C., and Bach, E.A. (2019). JAK/STAT signaling in stem cells and regeneration: from *Drosophila* to vertebrates. Development 146. 10.1242/dev.167643.

18. Galindo, M.I., Bishop, S.A., Greig, S., and Couso, J.P. (2002). Leg patterning driven by proximal-distal interactions and EGFR signaling. Science 297, 256–259. 10.1126/science.1072311.

19. Ishimaru, Y., Nakamura, T., Bando, T., Matsuoka, Y., Ohuchi, H., Noji, S., and Mito, T. (2015). Involvement of dachshund and Distal-less in distal pattern formation of the cricket leg during regeneration. Sci. Rep. 5, 8387. 10.1038/srep08387.

20. Nakamura, T., Mito, T., Tanaka, Y., Bando, T., Ohuchi, H., and Noji, S. (2007). Involvement of canonical Wnt/Wingless signaling in the determination of the positional values within the leg segment of the cricket *Gryllus bimaculatus*. Dev. Growth Differ. 49, 79–88. 10.1111/j.1440-169X.2007.00915.x.

21. Zeng, C., Liao, Q., Hu, Y., Shen, Y., Geng, F., and Chen, L. (2019). The Role of *Periplaneta americana* (Blattodea: Blattidae) in Modern Versus Traditional Chinese Medicine. J. Med. Entomol. 56, 1522–1526. 10.1093/jme/tjz081.

22. Ren, C., Chen, N., and Li, S. (2023). Harnessing “little mighty” cockroaches: Pest management and beneficial utilization. The Innovation 4. 10.1016/j.xinn.2023.100531.

23. Bando, T., Okumura, M., Bando, Y., Hagiwara, M., Hamada, Y., Ishimaru, Y., Mito, T., Kawaguchi, E., Inoue, T., Agata, K., et al. (2022). Toll signalling promotes blastema cell proliferation during cricket leg regeneration via insect macrophages. Development 149. 10.1242/dev.199916.

24. Mott, J.D., and Werb, Z. (2004). Regulation of matrix biology by matrix metalloproteinases. Curr. Opin. Cell Biol. 16, 558–564. 10.1016/j.ceb.2004.07.010.

25. Shapiro, S.D. (1998). Matrix metalloproteinase degradation of extracellular matrix: biological consequences. Curr. Opin. Cell Biol. 10, 602–608. 10.1016/s0955-0674(98)80035-5.

26. Upadhyay, A., Moss-Taylor, L., Kim, M.J., Ghosh, A.C., and O’Connor, M.B. (2017). TGF-β Family Signaling in *Drosophila*. Cold Spring Harb. Perspect. Biol. 9. 10.1101/cshperspect.a022152.

27. Kim, M., and McGinnis, W. (2011). Phosphorylation of Grainy head by ERK is essential for wound-dependent regeneration but not for development of an epidermal barrier. Proc. Natl. Acad. Sci. USA 108, 650–655. 10.1073/pnas.1016386108.

28. Raftery, L.A., and Sutherland, D.J. (1999). TGF-β family signal transduction in *Drosophila* development: from *Mad* to Smads. Dev. Biol. 210, 251–268. 10.1006/dbio.1999.9282.

29. Hao, I., Green, R.B., Dunaevsky, O., Lengyel, J.A., and Rauskolb, C. (2003). The *odd-skipped* family of zinc finger genes promotes *Drosophila* leg segmentation. Dev. Biol. 263, 282–295. 10.1016/j.ydbio.2003.07.011.

30. Brindley, H.H. (1897). On the Regeneration of the Legs in the Blattidœ. Proc. Zool. Soc. Lond 65, 903–916. 10.1111/j.1096-3642.1898.tb01392.x.

31. Campbell, G., and Tomlinson, A. (1995). Initiation of the proximodistal axis in insect legs. Development 121, 619–628. 10.1242/dev.121.3.619.

32. Li, L., Jing, A., Xie, M., Li, S., and Ren, C. (2021). Applications of RNA Interference in American Cockroach. J. Vis. Exp. 10.3791/63380.

33. Shirai, Y., Piulachs, M.D., Belles, X., and Daimon, T. (2022). DIPA-CRISPR is a simple and accessible method for insect gene editing. Cell Rep. Methods 2, 100215. 10.1016/j.crmeth.2022.100215.

34. Lin, G., and Slack, J.M. (2008). Requirement for Wnt and FGF signaling in *Xenopus* tadpole tail regeneration. Dev. Biol. 316, 323–335. 10.1016/j.ydbio.2008.01.032.

35. Taniguchi, Y., Watanabe, K., and Mochii, M. (2014). Notochord-derived hedgehog is essential for tail regeneration in *Xenopus* tadpole. BMC Dev. Biol. 14, 27. 10.1186/1471-213x-14-27.

36. Ho, D.M., and Whitman, M. (2008). TGF-β signaling is required for multiple processes during *Xenopus* tail regeneration. Dev. Biol. 315, 203–216. 10.1016/j.ydbio.2007.12.031.

37. Saito, N., Nishimura, K., Makanae, A., and Satoh, A. (2019). Fgf- and Bmp-signaling regulate gill regeneration in *Ambystoma mexicanum*. Dev. Biol. 452, 104–113. 10.1016/j.ydbio.2019.04.011.

38. Singh, B.N., Weaver, C.V., Garry, M.G., and Garry, D.J. (2018). Hedgehog and Wnt Signaling Pathways Regulate Tail Regeneration. Stem Cells Dev. 27, 1426–1437. 10.1089/scd.2018.0049.

39. Sun, G., and Irvine, K.D. (2014). Control of growth during regeneration. Curr. Top. Dev. Biol. 108, 95–120. 10.1016/b978-0-12-391498-9.00003-6.

40. Konstantinides, N., and Averof, M. (2014). A common cellular basis for muscle regeneration in arthropods and vertebrates. Science 343, 788–791. 10.1126/science.1243529.

41. Guarner, A., Manjón, C., Edwards, K., Steller, H., Suzanne, M., and Sánchez-Herrero, E. (2014). The *zinc finger homeodomain-2* gene of *Drosophila* controls *Notch* targets and regulates apoptosis in the tarsal segments. Dev. Biol. 385, 350–365. 10.1016/j.ydbio.2013.10.011.

42. Verghese, S., and Su, T.T. (2018). Ionizing radiation induces stem cell-like properties in a caspase-dependent manner in *Drosophila*. PLoS Genet. 14, e1007659. 10.1371/journal.pgen.1007659.

43. La Fortezza, M., Schenk, M., Cosolo, A., Kolybaba, A., Grass, I., and Classen, A.K. (2016). JAK/STAT signalling mediates cell survival in response to tissue stress. Development 143, 2907–2919. 10.1242/dev.132340.

44. Li, J., Wen, W., Zhang, S., Zhou, C., Feng, Y., and Li, X. (2022). The Expression and Function of lincRNA-154324 and the Adjoining Protein-Coding Gene *vmp1* in the Caudal Fin Regeneration of Zebrafish. Int. J. Mol. Sci. 23. 10.3390/ijms23168944.

45. Chang, J., Baker, J., and Wills, A. (2017). Transcriptional dynamics of tail regeneration in *Xenopus tropicalis*. Genesis 55. 10.1002/dvg.23015.

46. Arenas Gómez, C.M., Woodcock, R.M., Smith, J.J., Voss, R.S., and Delgado, J.P. (2018). Using transcriptomics to enable a plethodontid salamander (*Bolitoglossa ramosi*) for limb regeneration research. BMC Genomics 19, 704. 10.1186/s12864-018-5076-0.

47. Grotek, B., Wehner, D., and Weidinger, G. (2013). Notch signaling coordinates cellular proliferation with differentiation during zebrafish fin regeneration. Development 140, 1412–1423. 10.1242/dev.087452.

48. Raya, Á., Koth, C.M., Büscher, D., Kawakami, Y., Itoh, T., Raya, R.M., Sternik, G., Tsai, H.-J., Rodríguez-Esteban, C., and Izpisúa-Belmonte, J.C. (2003). Activation of Notch signaling pathway precedes heart regeneration in zebrafish. Proc. Natl. Acad. Sci. USA 100, 11889–11895. doi:10.1073/pnas.1834204100.

49. Beck, C.W., Christen, B., and Slack, J.M. (2003). Molecular pathways needed for regeneration of spinal cord and muscle in a vertebrate. Dev. Cell 5, 429–439. 10.1016/s1534-5807(03)00233-8.

50. Nakamura, K., and Chiba, C. (2007). Evidence for Notch signaling involvement in retinal regeneration of adult newt. Brain Res. 1136, 28–42. 10.1016/j.brainres.2006.12.032.

51. Blanco, E., Ruiz-Romero, M., Beltran, S., Bosch, M., Punset, A., Serras, F., and Corominas, M. (2010). Gene expression following induction of regeneration in *Drosophila* wing imaginal discs. Expression profile of regenerating wing discs. BMC Dev. Biol. 10, 94. 10.1186/1471-213x-10-94.

52. Liu, W., Singh, S.R., and Hou, S.X. (2010). JAK-STAT is restrained by Notch to control cell proliferation of the *Drosophila* intestinal stem cells. J. Cell Biochem. 109, 992–999. 10.1002/jcb.22482.

53. Vallecillo-García, P., Orgeur, M., Vom Hofe-Schneider, S., Stumm, J., Kappert, V., Ibrahim, D.M., Börno, S.T., Hayashi, S., Relaix, F., Hildebrandt, K., et al. (2017). Odd skipped-related 1 identifies a population of embryonic fibro-adipogenic progenitors regulating myogenesis during limb development. Nat. Commun. 8, 1218. 10.1038/s41467-017-01120-3.

54. Stumm, J., Vallecillo-García, P., Vom Hofe-Schneider, S., Ollitrault, D., Schrewe, H., Economides, A.N., Marazzi, G., Sassoon, D.A., and Stricker, S. (2018). Odd skipped-related 1 (Osr1) identifies muscle-interstitial fibro-adipogenic progenitors (FAPs) activated by acute injury. Stem Cell Res. 32, 8–16. 10.1016/j.scr.2018.08.010.

55. de Celis, J.F., Tyler, D.M., de Celis, J., and Bray, S.J. (1998). Notch signalling mediates segmentation of the *Drosophila* leg. Development 125, 4617–4626. 10.1242/dev.125.23.4617.

56. Godt, D., Couderc, J.L., Cramton, S.E., and Laski, F.A. (1993). Pattern formation in the limbs of *Drosophila*: *bric à brac* is expressed in both a gradient and a wave-like pattern and is required for specification and proper segmentation of the tarsus. Development 119, 799–812. 10.1242/dev.119.3.799.

57. Bosch, M., Bishop, S.A., Baguña, J., and Couso, J.P. (2010). Leg regeneration in *Drosophila* abridges the normal developmental program. Int. J. Dev. Biol. 54, 1241–1250. 10.1387/ijdb.093010mb.

58. Sinigaglia, C., Almazán, A., Lebel, M., Sémon, M., Gillet, B., Hughes, S., Edsinger, E., Averof, M., and Paris, M. (2022). Distinct gene expression dynamics in developing and regenerating crustacean limbs. Proc. Natl. Acad. Sci. USA 119, e2119297119. 10.1073/pnas.2119297119.

59. Stappenbeck, T.S., and Miyoshi, H. (2009). The role of stromal stem cells in tissue regeneration and wound repair. Science 324, 1666–1669. 10.1126/science.1172687.

60. Kim, D., Paggi, J.M., Park, C., Bennett, C., and Salzberg, S.L. (2019). Graph-based genome alignment and genotyping with HISAT2 and HISAT-genotype. Nat. Biotechnol. 37, 907–915. 10.1038/s41587-019-0201-4.

61. Love, M.I., Huber, W., and Anders, S. (2014). Moderated estimation of fold change and dispersion for RNA-seq data with DESeq2. Genome Biol. 15, 550. 10.1186/s13059-014-0550-8.

62. Ligges, U., and Maechler, M. (2003). scatterplot3d - An R Package for Visualizing Multivariate Data. J. Stat. Softw. 8, 1–20. 10.18637/jss.v008.i11.

63. Yu, G., Wang, L.G., Han, Y., and He, Q.Y. (2012). clusterProfiler: an R package for comparing biological themes among gene clusters. Omics 16, 284–287. 10.1089/omi.2011.0118.

64. Langmead, B., and Salzberg, S.L. (2012). Fast gapped-read alignment with Bowtie 2. Nat. Methods 9, 357–359. 10.1038/nmeth.1923.

65. Zhang, Y., Liu, T., Meyer, C.A., Eeckhoute, J., Johnson, D.S., Bernstein, B.E., Nusbaum, C., Myers, R.M., Brown, M., Li, W., and Liu, X.S. (2008). Model-based analysis of ChIP-Seq (MACS). Genome Biol. 9, R137. 10.1186/gb-2008-9-9-r137.

66. Yu, G., Wang, L.G., and He, Q.Y. (2015). ChIPseeker: an R/Bioconductor package for ChIP peak annotation, comparison and visualization. Bioinformatics 31, 2382–2383. 10.1093/bioinformatics/btv145.

67. Kechin, A., Boyarskikh, U., Kel, A., and Filipenko, M. (2017). cutPrimers: A New Tool for Accurate Cutting of Primers from Reads of Targeted Next Generation Sequencing. J. Comput. Biol. 24, 1138–1143. 10.1089/cmb.2017.0096.

68. Thorvaldsdóttir, H., Robinson, J.T., and Mesirov, J.P. (2013). Integrative Genomics Viewer (IGV): high-performance genomics data visualization and exploration. Brief Bioinform. 14, 178–192. 10.1093/bib/bbs017.

