## Supplemental figures for "Two transcriptional cascades orchestrate cockroach leg regeneration"

**This PDF file includes:**

Figs. S1 to S7

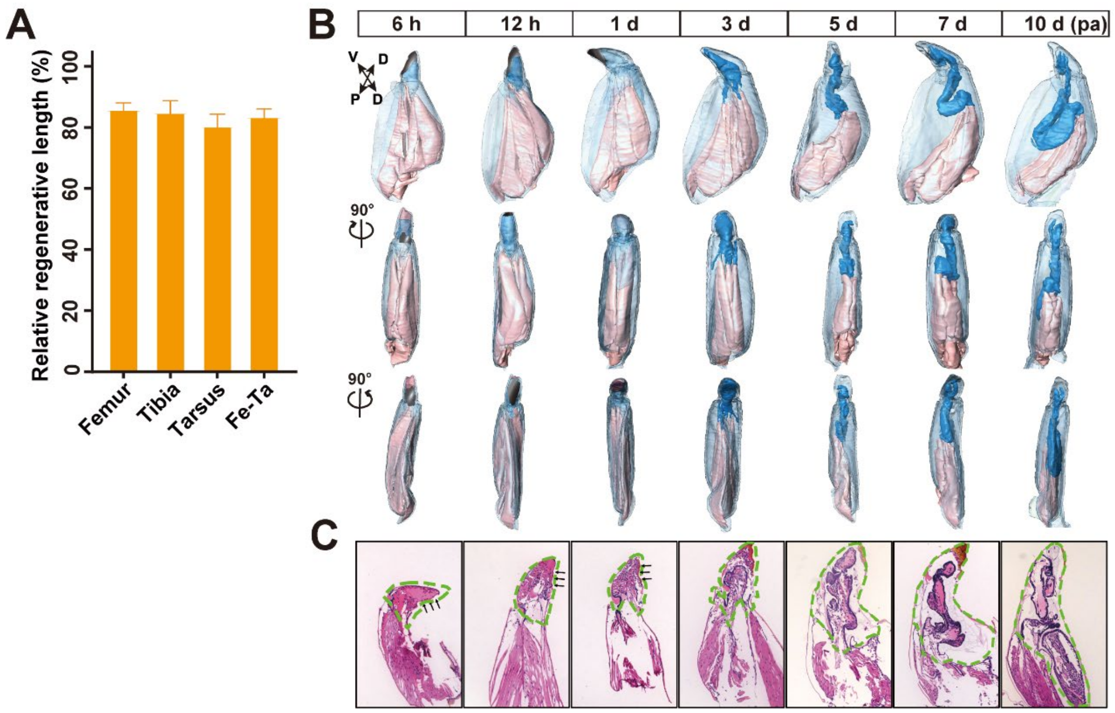


**Fig. S1**. **Measurement of the regenerative length and morphological profiles of leg regeneration in American cockroach. Related to Fig. 1.** (A). Measurement of the relative regenerative length after amputation at trochanter-femur joint site. The relative regenerative length of femur, tibia and tarsus are measured separately and together. (B). 3D model of the morphological profiles of regenerating legs made by μ-CT. The other three directions were adopted to show the regenerating microstructural objects. (C). HE staining of regenerating legs under transection, the arrows indicate wound sites and green dotted lines cover the regenerative regions.

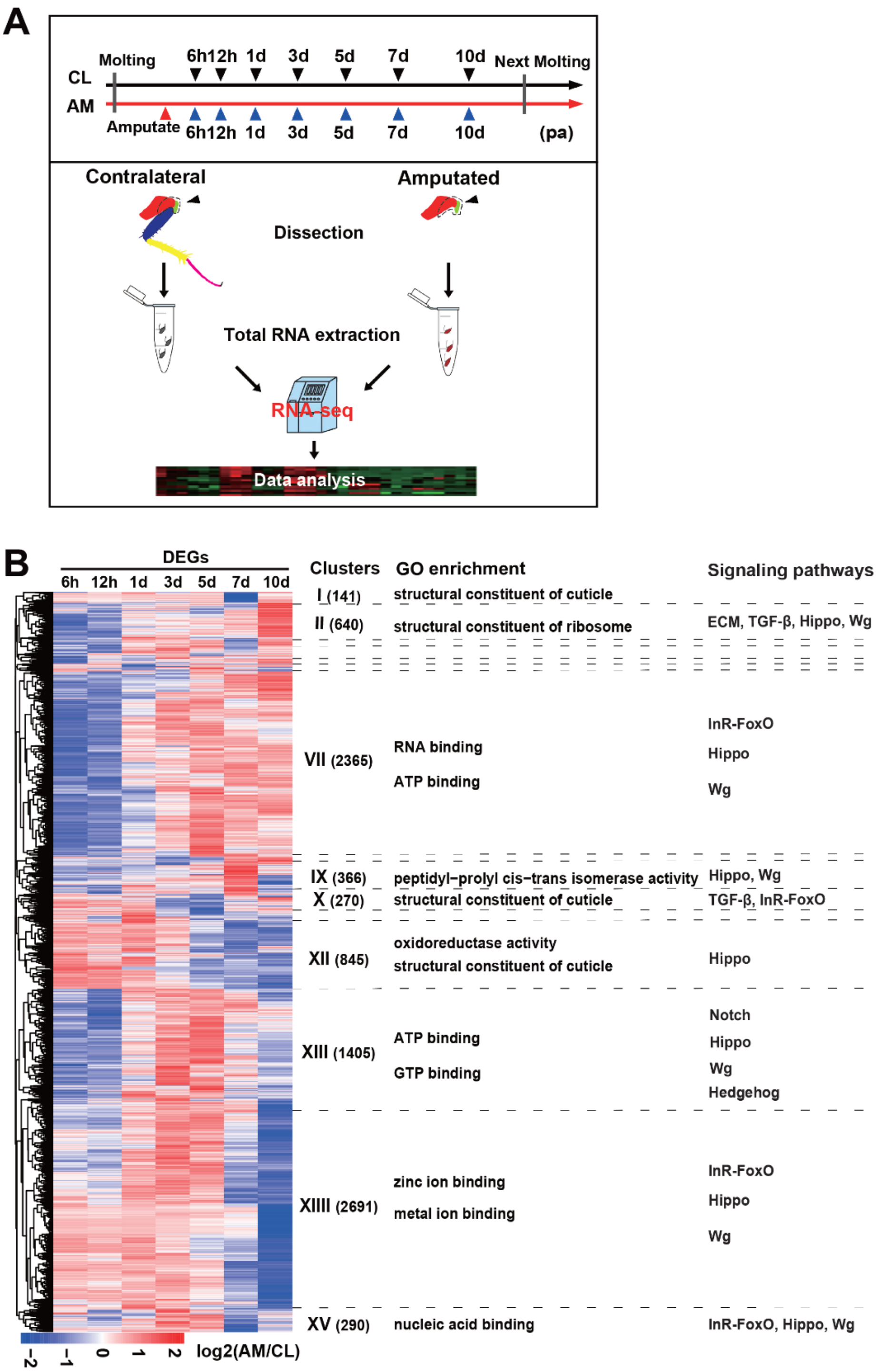


**Fig. S2. Transcriptome sequencing strategy and enrichment analyses. Related to Fig. 1.** (A). Schematic representation of the strategy to prepare the samples for RNA-Seq at 7 time points during developmental/homeostatic (CL group) and regenerative (AM group) stages. The tissues within the dashed line region were collected for sequencing. (B). Heatmap showing k-means clustering (k = 15) of 9,473 dynamically expressed genes across regeneration. Each horizontal line describes the relative expression of a single gene. Cluster numbers are indicated using Roman numerals and the number of genes in each cluster are added. Some clusters with fewer gene numbers are not shown. Enriched processes from GO and signaling pathways within the main clusters are displayed in the right panels.

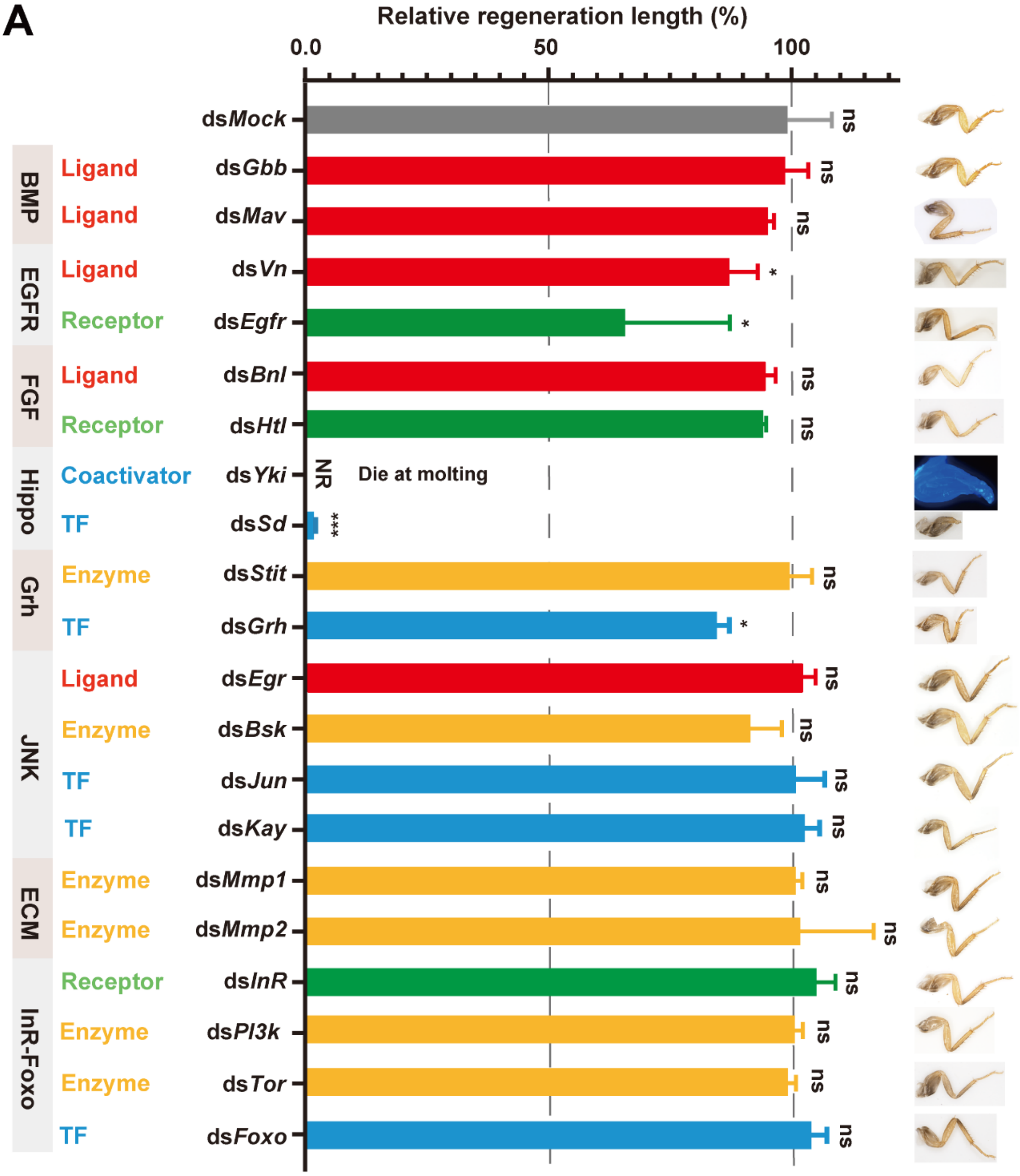


**Fig. S3**. **Screening of signaling pathways necessary for leg regeneration. Related to Fig. 2.** Eight signaling or pathways were selected and the key components ligands, receptors, coactivators, enzymes, and TFs were labelled in different colors. The legs of the insects in the *Yki* group were dissected and stained with DAPI because they encountered a molting barrier, NR: no visible regenerated leg was observed. The significance of differences for regeneration abilities were analyzed by two-tailed Student's *t test*. *: $P$ <0.05, **: $P$ <0.01, ***: $P$ <0.001, "ns" stands for "no significant difference". n=3.

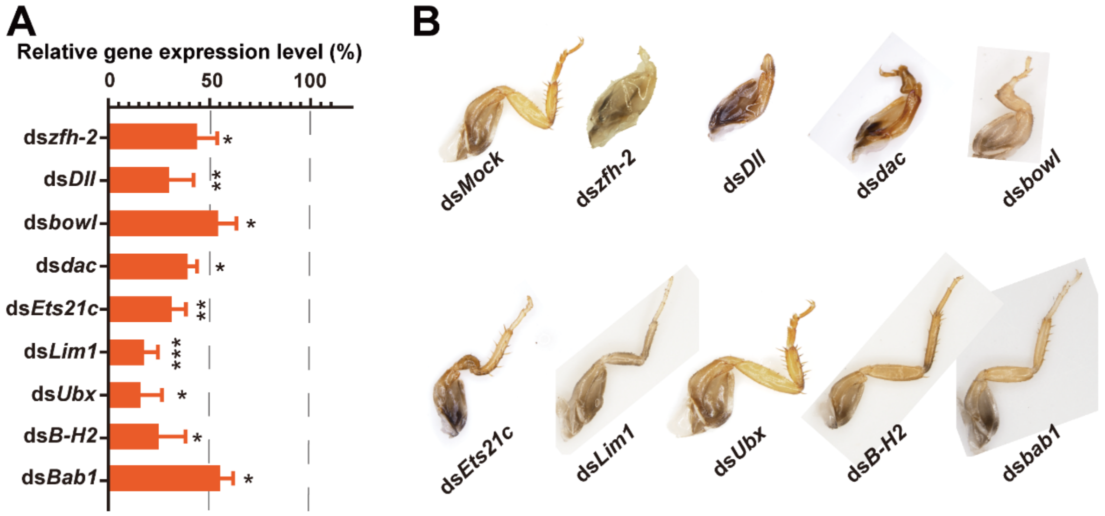


**Fig. S4**. **RNAi efficiency detection and phenotypes of TFs. Related to Fig. 3.** (A). The RNAi efficiency of these nine TFs were detected by qRT–PCR. The significance of differences for regeneration abilities were analyzed by two-tailed Student's *t test*. *: $P < 0.05$, **: $P < 0.01$, ***: $P < 0.001$. n=3. (B). The original and big sized photos of leg phenotypes related to fig. 3D.

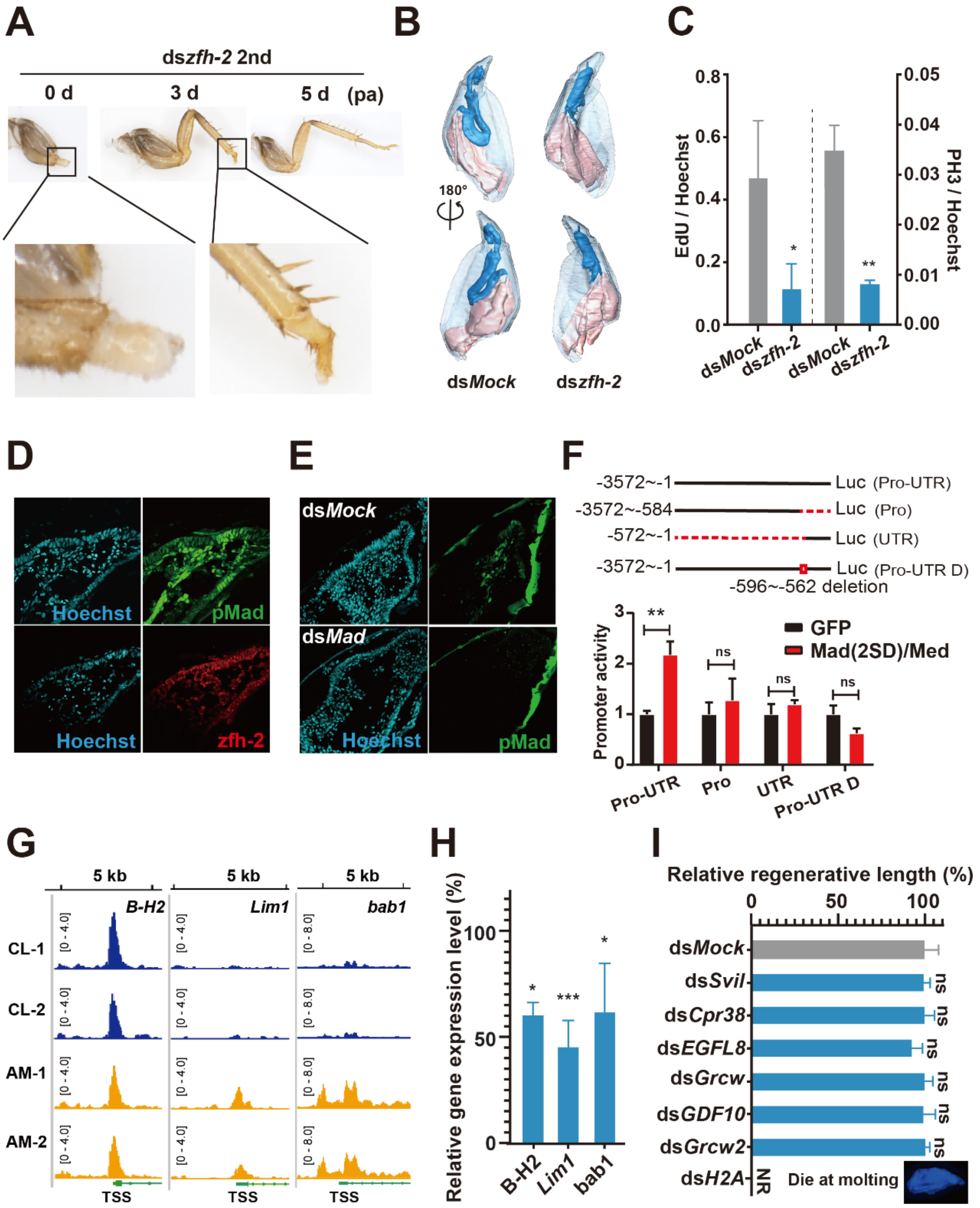


**Fig. S5**. **zfh-2 contributes to blastema proliferation and morphogenesis under the control of BMP and JAK-STAT signaling pathways**. **Related to Fig. 4.** (A). Phenotype of regenerated legs when the second design of ds*zfh-2* was injected at 0, 3 and 5 dpa. n=3. (B). Structure of regenerated leg at 10 dpa under ds*zfh-2* treatment detected by μ-CT. (C). The statistics of the percentages of EdU and PH3 positive cells. dsRNAs were injected at 0 dpa and the samples were harvested at 3 dpa, the EdU was injected at 6 h before harvest. n=3. (D). The colocalization of pMad and zfh-2 proteins in regenerative cells at 3 dpa were stained using serial tissue section method. (E). Antibody-based immunostaining of pMad protein level in ds*Mock* and ds*Mad* treatment groups at 3 dpa, n=3. (F). Dual luciferase assay for the detection of Mad(2SD)/Med binding activity on the truncated and deleted promoter of *zfh-2*. Pro: promoter; UTR: 5′ untranslated region; Pro-UTR-D: promoter and 5′ untranslated region with −596-−562 deletion. (G).

Chromatin accessibility around the TSS region of *B-H2 Lim1,* and *bab1* were detected using ATAC-Seq. (H). The relative expression of B-H2, Lim1, and bab1 when they were knocked down simultaneously. (I). Measurement of relative regenerative length of regenerated legs under dsRNA treatments. n=3. The significance of differences was analyzed by two-tailed Student's *t test*. **: $P$ <0.01, ***: $P$ <0.001, "ns" stands for "no significant difference", NR: no visible regenerated leg was observed.

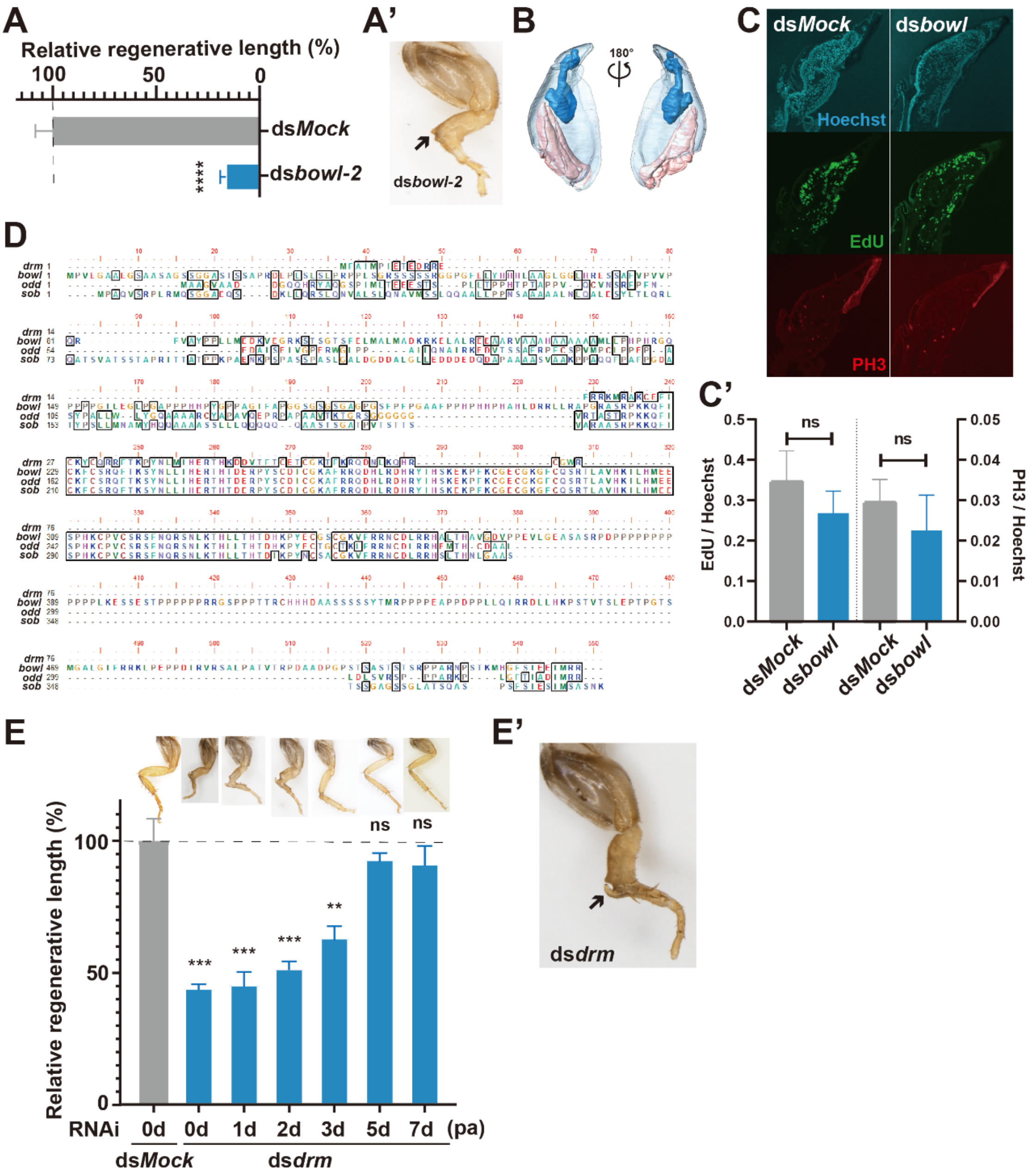


**Fig. S6. The odd-skipped family members bowl and drm regulate leg morphogenesis. Related to Fig. 5.** (A-A'). Phenotype and relative regenerative length of regenerated legs when the second design of ds*bowl* was injected. n=3. (B). Structure of regenerated leg at 10 dpa under ds*bowl* treatment detected by µ-CT. (C-C'). EdU and PH3 staining of proliferation positive cells. dsRNAs were injected at 0 dpa and the samples were harvested at 3 dpa, the EdU was injected at 6 h before harvest. The rates of proliferation positive cells were calculated by EdU/Hoechst and PH3/Hoechst. (D). Alignment of the amino acids of odd-skipped family proteins. (E-E'). Phenotypes and relative regenerative length of regenerated legs when *drm* was knocked down at six different time points post amputation (E). The close-up of regenerated leg when *drm* was knocked down at 0 dpa (E'). The significance of differences was analyzed by two-tailed Student's *t test*. **: *P* <0.01, ***: *P* <0.001, "ns" stands for "no significant difference".

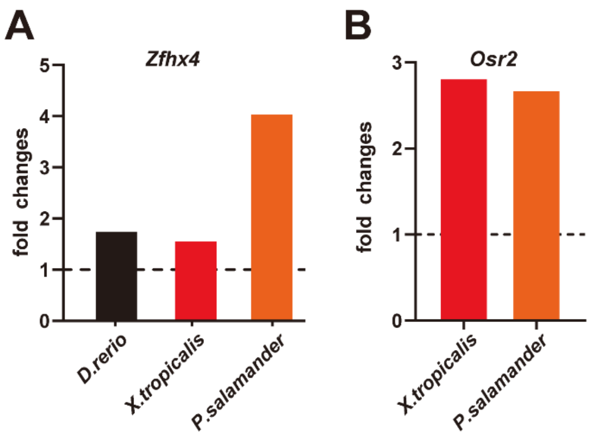


**Fig. S7**. **Relative expression of *zfh-2* ortholog *Zfhx4* and *bowl* ortholog *Osr2* in vertebrates. Related to Discussion.** (A). Fold changes of *Zfhx4* in caudal fin regeneration of *D. rerio*, tail regeneration of *X. tropicalis*, and limb regeneration of *P. salamander*. (B). Up regulated fold changes of *Osr2* in tail regeneration of *X. tropicalis* and limb regeneration of *P. salamander*.
